# Cotranscriptional RNA strand exchange underlies the gene regulation mechanism in a purine-sensing transcriptional riboswitch

**DOI:** 10.1101/2021.10.25.465737

**Authors:** Luyi Cheng, Elise N. White, Naomi L. Brandt, Angela M Yu, Alan A. Chen, Julius B. Lucks

## Abstract

RNA folds cotranscriptionally to traverse out-of-equilibrium intermediate structures that are important for RNA function in the context of gene regulation. To investigate this process, here we study the structure and function of the *Bacillus subtilis yxjA* purine riboswitch, a transcriptional riboswitch that downregulates a nucleoside transporter in response to binding guanine. Although the aptamer and expression platform domain sequences of the *yxjA* riboswitch do not completely overlap, we hypothesized that a strand exchange process triggers its structural switching in response to ligand binding. *In vivo* fluorescence assays, structural chemical probing data, and experimentally informed secondary structure modeling suggest the presence of a nascent intermediate central helix. The formation of this central helix in the absence of ligand appears to compete with both the aptamer’s P1 helix and the expression platform’s transcriptional terminator. All-atom molecular dynamics simulations support the hypothesis that ligand binding stabilizes the aptamer P1 helix against central helix strand invasion, thus allowing the terminator to form. These results present a potential model mechanism to explain how ligand binding can induce downstream conformational changes by influencing local strand displacement processes of intermediate folds that could be at play in multiple riboswitch classes.

## INTRODUCTION

RNA folds during transcription (1-3). During this cotranscriptional folding process, RNA molecules traverse sequential structural transitions that lead to the formation of helices, loops, junctions, pseudoknots, and other elements that are important for RNA function (4, 5). Cotranscriptional RNA folding has been shown to be important for establishing the functional folds of catalytic RNAs (6, 7), assembling the ribosome and spliceosome (8-10), regulating RNA processing (11), and facilitating gene regulation (12). While we are beginning to understand the importance of cotranscriptional RNA folding, we still lack a complete mechanistic understanding of how folding pathways are encoded within specific RNA sequences, and the details of how these folding processes occur within a range of cellular functions (13).

Riboswitches have emerged as important model systems for uncovering principles of cotranscriptional folding (14, 15). These compact non-coding RNA elements regulate gene expression in *cis* in response to the binding of a specific chemical ligand, thus acting as RNA biosensors in a wide distribution of bacteria species as well as fungi (16-18). Riboswitches achieve this function through their architecture which consists of a ligand binding aptamer domain (AD) that lies upstream of an expression platform (EP) whose folding state determines the functional gene expression outcome (19). Riboswitches are typically classified by the cognate ligand they respond to, the type of regulatory control the EP exerts on either the translational, transcriptional, or RNA splicing level, and whether they upregulate or downregulate gene expression in response to ligand binding (17, 20). Transcriptional riboswitches contain an EP that includes an intrinsic terminator hairpin that, once folded, prevents transcription elongation (19). These riboswitches are particularly important model systems for understanding principles of cotranscriptional folding because AD folding, ligand binding, and EP folding need to occur within the short time window of active transcription (21).

While studies have extensively characterized the structures of riboswitch ADs (22-26), current research is only just beginning to understand how structural features induced by ligand binding in the AD can alter the EP structure and determine gene expression outcomes during transcription (27-33). Extensive progress has come from the study of transcriptional ‘ON’ riboswitches that contain an overlap in sequence between the AD and EP that lends to mutually exclusive folds of these two domains. In these cases, studies have shown that ligand-binding stabilizes the AD fold, which prevents the EP from forming a terminator hairpin (27-29, 34). On the other hand, when a ligand is not bound, EP folding disrupts the AD structure through RNA strand displacement. During this process, the mutually exclusive helices interconvert — a set of nucleating base pairs seed the formation of a new helix while unwinding the existing helix (35, 36). Thus, when there is an overlap in sequence between the ligand binding portion of the AD and EP, there appears to be a clear mechanism by which ligand binding can affect downstream folding of the EP through RNA strand displacement (37, 38).

However, out of all identified riboswitch classes to date, many of them do not encode a direct sequence overlap between the AD and EP (20, 39, 40). This observation raises an important question for understanding the mechanisms of these riboswitches: how can ligand binding in the AD “communicate” downstream structural changes and determine folding outcomes of the EP (37, 38)? Given the role of strand displacement mechanisms among diverse riboswitches (29, 40, 41), as well as within the cotranscriptional folding pathways of other important non-coding RNAs such as the signal recognition particle (SRP) RNA (42), we hypothesize that riboswitches without direct AD and EP overlap still rely on a ligand-dependent strand exchange process to differentiate their folded states and enact regulatory function, however with an unknown mechanism to inhibit strand exchange in the presence of ligand.

To investigate this hypothesis, we focused our studies on the *Bacillus subtilis yxjA* guanine riboswitch, a transcriptional ‘OFF’ riboswitch that downregulates the expression of a nucleoside transporter protein in response to excess guanine as a part of nucleotide biosynthesis pathway regulation (43, 44). As part of the purine-sensing riboswitch family, the *yxjA* riboswitch shares 75.4% sequence conservation and a similar predicted AD structure with the adenine-responsive *Bacillus subtilis pbuE* riboswitch. These AD aptamers consist of a P1 helix at the base of a three-way junction (3WJ), with two protruding P2 and P3 helices that are oriented through a kissing hairpin pseudoknot interaction (45-47) (Figure 1A). Within the ligand binding pocket provided by the 3WJ, the J2/3 junction sequence forms a triple helix with the top of P1 while a single nucleotide in the J3/1 junction sequence determines the ligand binding specificity. Despite the similarities between the *yxjA* and *pbuE* ADs however, important differences remain, including the observation that the *pbuE* riboswitch is an ‘ON’ regulator with inverse expression logic to the *yxjA* riboswitch. The *pbuE* riboswitch EP also shares direct sequence overlap with a large portion of the AD, extending through the P3 helix, which explains how ligand binding can dictate which mutually exclusive structure to adopt. In contrast, the *yxjA* AD core 3WJ structure and the predicted EP fold appear to be compatible with each other at a secondary structural level (Figure 1C and D). Thus, *yxjA* is a model riboswitch for investigating how ligand binding may inhibit cotranscriptional RNA strand displacement processes that help form of downstream RNA structures.

**Figure 1.**
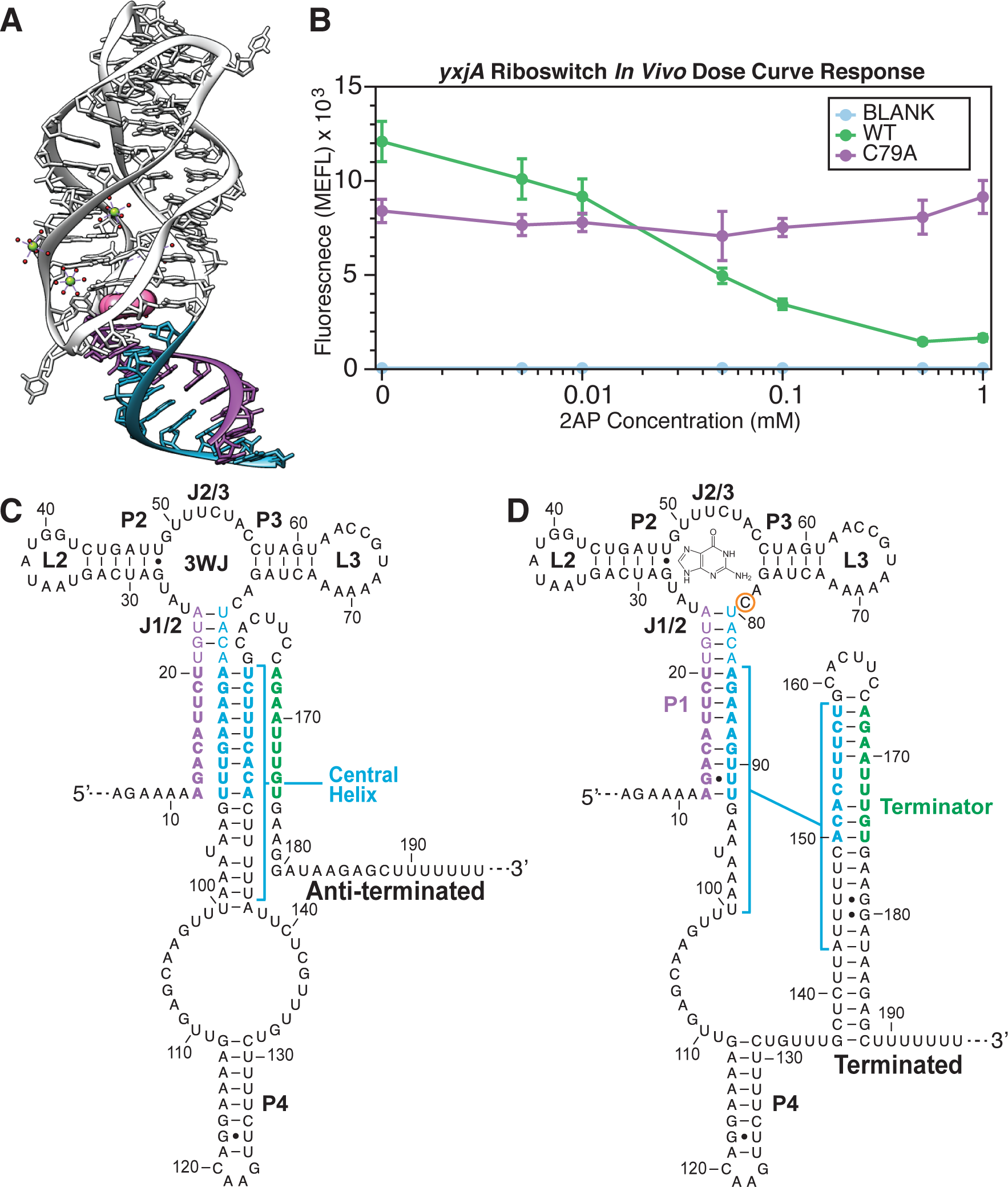
Structure schematic and ligand-dependent functional response of the *yxjA* riboswitch. **(A)** Color coded crystal structure (PDB ID 1Y27) of the purine riboswitch aptamer in the ligand-bound holo conformation. The ligand is colored pink, the 5′ side of the P1 helix purple, and the 3′ side of the P1 helix teal (53). **(B)** Dose response curve of the *yxjA* riboswitch regulation in *E. coli* cells. *E. coli* cells were transformed with plasmids containing riboswitch expression constructs and characterized for SFGFP fluorescence using flow cytometry after 5 hours of growth in M9 minimal media subcultures with or without 2-aminopurine (2AP) added. A blank control with a constitutive promoter and no SFGFP was used to measure any cellular background fluorescence levels in response to ligand in the media. A non-ligand-responsive riboswitch construct (C79A) was used as a negative control to account for non-riboswitch regulated changes in fluorescence in response to ligand in the media. WT represents the wild type *yxjA* riboswitch sequence in the expression construct. Riboswitch secondary structure models in **(C)** represent the predicted anti-terminated conformation consisting of an intact aptamer domain, a partially disrupted P1 stem, and the central helix formed. Secondary structure models in **(D)** represent the predicted terminated conformation consisting of the intact aptamer domain, a fully formed P1 helix and the formed intrinsic terminator expression platform. Secondary structure models are based on minimum free energy predictions from Muhlbacher and Lafontaine, 2007 (43). Nucleotide regions are colored to represent base pairs that are mutually exclusive between the anti-terminated and terminated structure models (teal), nucleotides that are predicted to complete the P1 helix on the 5′ side (purple), and nucleotides predicted to complete the terminator on the 3′ side (green). In (D), the ligand is shown in proximity to the 3WJ ligand-binding pocket, where the residue C79 that makes ligand contacts is circled. Dots in (B) represent averages from three biological replicates, each performed in triplicate technical replicates for a total of nine data points (n=9), with error bars representing standard deviation.

In this work, we present a series of functional gene expression studies of rationally designed *yxjA* riboswitch mutants, motivated by preliminary RNA structure predictions, to propose a mechanism for the *yxjA* riboswitch tò serve as a model for several identified riboswitch classes. We conducted SHAPE-Seq structure probing experiments of two intermediate and the full length riboswitch sequences within stalled transcription elongation complexes and under equilibrium refolded conditions to generate reactivity data, which we then used as restraints for RNA structure prediction algorithms. The resulting experimentally informed structure models revealed an intermediate central helix structure that is mutually exclusive with the bottom of the P1 stem and the terminator hairpin of the EP. Mutants that disrupt base pairing of this central helix break regulatory function, while rescue mutants restore function, demonstrating its importance and suggesting that ligand binding in the AD structurally regulates the formation of the central helix. To investigate this further, we performed restrained all-atom molecular dynamics (MD) simulations that demonstrated ligand binding helps to stabilize the P1 helix against strand invasion by the central helix, thereby providing a mechanism to explain how ligand binding in the AD can influence folding of the downstream EP.

Based on our results, we propose that the RNA strand exchange plays a critical role in the *yxjA* riboswitch mechanism and that the mechanism may rely on a competition between two strand exchange processes – central helix formation through unzipping the base of the P1 stem, and terminator formation through unzipping of the central helix – with the winner of the competition determining the riboswitch folded state and regulatory outcome. Furthermore, we suggest that ligand binding can bias this strand exchange process by stabilizing the P1 helix, providing a larger energetic barrier for strand displacement by the central helix and thereby allowing the terminator to form more efficiently to make an ‘OFF’ regulatory decision. We believe this work adds valuable knowledge for understanding the mechanism of how riboswitches with modular or non-overlapping aptamer domains and expression platforms can make genetic decisions and provides a possible overarching mechanism for a broad number of riboswitch classes. We anticipate that competing RNA strand displacement processes may play a role in other cotranscriptional RNA folding phenomena, and that dissecting riboswitch mechanisms can be ideal for designing synthetic riboswitch systems.

## MATERIALS AND METHODS

### Cloning and plasmid construction

To prepare each riboswitch construct, the *Bacillus subtilus yxjA* (nupG) 5′-untranslated region (UTR) sequence encoding the riboswitch as wildtype (WT) and mutants was cloned into p15a plasmid backbones with chloramphenicol resistance. Inverse polymerase chain reaction (iPCR) was used to introduce the riboswitch sequence downstream of the *Escherichia coli* sigma 70 consensus promoter (denoted J23119 in the registry of standardized biological parts) and upstream of a ribosome binding site and the coding sequence of the superfolder green fluorescent protein (SFGFP) reporter gene. Various lengths of the riboswitch leader sequence from the *B. subtilis* genome were similarly inserted between the promoter and riboswitch sequence when optimizing constructs for testing functional response to ligand binding. For creating different mutants using iPCR, primers were ordered from Integrated DNA Technologies (IDT) and phosphorylated using a New England Biosciences (NEB) phosphorylation kit (PNK enzyme, 10X T4 DNA Ligase Buffer, primers). 100 µL PCR reactions were mixed with 1 – 10 ng/µL of DNA template, 200 µM dNTPs, 1X Phusion Buffer, 400 nM each of phosphorylated forward and reverse primers and 0.25 µL of Phusion Polymerase (2,000 U/mL) (NEB). PCR products were run on 1% agarose gels to confirm successful amplification. DpnI (NEB) was used for digestion and T4 DNA Ligase (NEB) for ligation. Final amplified products were transformed into NEBTurbo competent cells and plated on chloramphenicol agar plates and incubated overnight at 37°C. Single isolated colonies were selected the next day to inoculate cultures in LB media, mini-prepped, and then sequence confirmed using Sanger Sequencing (Quintara).

### *In vivo* flow cytometry fluorescence reporter assay

Fluorescence assays (Supplementary Figure S1) on the flow cytometer were performed in *E. coli* strain TG1 cells with three independent biological replicates. Plasmids tested for each replicate, including *yxjA* WT (C79T mutation), *yxjA* ON mutant control (C79A mutation), *yxjA* experimental mutants, empty control plasmid (pJBL001), and constitutively expressed SFGFP control plasmid (pPDC1095), were transformed into competent *E. coli* TG1 cells, plated on LB agar plates containing 34 mg/mL chloramphenicol, and incubated overnight at 37°C, for approximately 18 hours. After overnight incubation, the plates containing transformed *E. coli* colonies were removed from the incubator and kept at room temperature at benchtop for approximately 5 hours. The three replicates of riboswitch constructs and controls were picked from plate colonies to inoculate 300 µL of LB media containing 34 mg/mL chloramphenicol in a 2 mL 96-well block. The block was covered with a breathable seal and incubated in a benchtop shaker at 37°C overnight while shaking at 1000 rpm for 18 hours. The next day, cultures for each biological replicate were used to inoculate subcultures in triplicate for each of three technical replicates. Each technical replicate was created by transferring 4 µL of an overnight shaking culture to each of two 96-well blocks containing 196 µL of M9 media (1X M9 salts, 1 mM thiamine hydrochloride, 0.4% glycerol, 0.2% casamino acids, 2 mM MgSO_4_, 0.1 mM CaCl_2_, 34 mg/mL chloramphenicol) with either 1 mM 2-aminopurine (2AP) dissolved in 1% acetic acid (with ligand condition) or an equal volume of 1% acetic acid (no ligand condition). The block was covered with a Breath-Easier sealing membrane (Sigma-Aldrich) and incubated in a benchtop shaker at 37°C while shaking at 1000 rpm for 5 hours unless otherwise indicated. Samples for fluorescence analysis were prepared after incubation by transferring 4 µL of exponential phase culture to 196 µL of phosphate buffered saline (1X PBS) and kanamycin (50 ug/mL) in 96-well v-bottom plates (Costar).

Fluorescence measurements were collected on a BD Accuri C6 Plus with a CSampler Plus with the BD CSampler Plus software. *E. coli* events were collected using the FSC-A threshold set to 2,000. GFP fluorescence was measured by collecting FITC signal on the FL1-A channel.

Analysis was carried out by using the FlowJo software (v10.6.1) (Supplementary Figure S5). An ellipsoid gate was drawn around the *E. coli* cell population on the FSC-A vs SSC-A profile. Corresponding arbitrary fluorescence units collected by the flow cytometer were plotted against known standard molecules of equivalent fluorescein (MEFL) values from Spherotech rainbow 8-peak calibration beads (Lot: 8137522). The calibration curve was created by calculating the linear regression between measured relative fluorescence units (RFU) of each bead peak with the manufactured supplied MEFL value for each peak, with the y-intercept set to zero. The mean fluorescence of peaks in the FITC channel corresponding to the gated *E. coli* cell population was converted to a MEFL value by multiplying by the slope of the calibration curve.

### *In vivo* bulk fluorescence reporter assay

Samples for bulk fluorescence assays (Supplementary Figure S2 and S3) on the plate reader were prepared by transferring 50 µL of culture from the procedure above for flow cytometry assays to 50 µL of phosphate buffered saline (1X PBS) for bulk fluorescence measurements with a Biotek Synergy H1 plate reader and gain set to 70. Analysis was carried out by subtracting blank optical density (OD) measurements and fluorescence values of M9/PBS buffer mix from each sample containing cells. Arbitrary fluorescence units for each well were divided by OD_600_ measurements in that well to normalize for the amount of cells in each sample, prior to performing averages and standard deviations across replicates. For the time course optimization experiment, measured samples were successively collected from the same subculture sampled.

### DNA template preparation

DNA template libraries for cotranscriptional and equilibrium SHAPE-Seq experiments (Supplementary Table S2) were prepared by PCR amplifying plasmids containing the *yxjA* riboswitch sequence. 500 µL PCR reactions were mixed to a final concentration of 1X ThermoPol Buffer, 250 nM each of forward and reverse primers (Supplementary Table S3), 1.25 mM dNTPs, 5 uL of Taq Polymerase (5,000 U/mL) (NEB), and 100 – 300 ng of DNA template. Templates were amplified for 30 cycles and then purified on a 1% agarose gel. Bands were visualized using UV and then extracted using a Qiagen Gel Extraction kit and concentrations were measured on a Qubit.

### *In vitro* transcription assay with SHAPE modification

50 uL reaction mixtures contained 5 nM linear DNA template for each of the template lengths tested (Supplementary Table S2), 1 U of *E. coli* RNAP holoenzyme (New England BioLabs), transcription buffer (20 mM Tris-HCl, pH 8.0, 0.1 mM EDTA, 1 mM DTT, and 50 mM KCl), 0.1 mg/mL bovine serum albumin (New England BioLabs), 200 uM ATP, GTP, CTP, and UTP. Reactions also contained either 1 mM final concentration of 2AP dissolved in 1% acetic acid (with ligand condition) or the equivalent volume of 1% acetic acid (no ligand condition). The reaction tube was incubated for 10 minutes at 37°C to form open transcription complexes. Transcription was initiated with the addition of MgCl_2_ at a final concentration of 5 mM and rifampicin at 10 µg/mL. Reactions were incubated for another 30 seconds at 37°C. Each 50 µL was then split in half with 25 uL pipetted into a tube containing 2.78 µL of 400 mM benzoyl cyanide (BzCN) dissolved in DMSO (final concentration 40 mM) for the SHAPE reagent probed (+) channel and the other 25 µL pipetted into a tube containing an equivalent volume of DMSO for the negative control (-) channel. 75 µL of TRIzol (Invitrogen) was then pipetted into the tube to stop transcription.

For RNA extraction and purification, after adding 20 µL of chloroform, 70 µL of the aqueous phase was extracted and then ethanol precipitated with the addition of 50 µL of isopropanol (following TRIzol extraction protocol) and rested for 10 minutes at room temperature. Samples were spun down for 10 minutes at 4°C into a pellet, then aspirated and washed with 500 µL of 70% ethanol. Dried pellets were resuspended in 43 µL of water and incubated with 5 µL of 10X TURBO DNase buffer and 2 µL of TURBO DNase (Thermo Fisher) for 1 hour at 37°C. After digestion, 150 µL of TRIzol and 40 µL of chloroform were added for another round of extraction and purification. 140 µL of the aqueous phase was extracted then ethanol precipitated with the addition of 100 µL of isopropanol and following the same steps as above.

### Cotranscriptional SHAPE-Seq library preparation

Extracted RNA from IVT and SHAPE modification was used to prepare SHAPE-Seq libraries with the methods described in Watters, et al., 2016 and Yu, et al., 2021 (27, 42). Additional oligonucleotides (IDT) used for library preparation can be found in Supplementary Table S3. Several modifications to previously published protocols used in this work are as follows:

Linker ligation: 0.2 pmol of RNA linker was added.

Adapter ligation: The DNA adapter sequence and ligation protocol was adapted from the Structure-seq2 protocol described in Ritchey, et al., 2017 (48). Reverse transcribed cDNA product was resuspended in 7 µL of water and mixed with 0.5 µL of 100 µM of dumbbell adapter (final concentration 50 pmol). Reactions were incubated at 95°C for 2 minutes followed by incubation at room temperature for 3 minutes. The ligation reaction was then set up by combining the cDNA and adapter mixture with 1X T4 DNA Ligase Buffer (NEB), 500 mM Betaine, 20% PEG-8000, and 2.5 µL T4 DNA Ligase in a 25 µL volume solution. The ligation reaction was incubated on a thermocycler at 16°C for 6 hours, 30°C for 6 hours, 65°C for 15 minutes, then 4°C until ready for ethanol precipitation. Subsequent steps were performed according to previously published protocols.

For sequencing library preparation, the average DNA length and quality of each library sample was determined by running samples on an 8% non-denaturing PAGE gel stained with SYBR Gold. Each library sample concentration was measured on a Qubit 3.0 and pools were balanced with each library at equimolar amounts. Pooled libraries also included an 8% PhiX spike-in to increase sequence diversity. Sequencing data was then obtained by sequencing the pooled library samples on an Illumina MiSeq using the v3 MiSeq Reagent Kit and 2 x 75 bp paired-end reads. Reactivity analyses using FASTQ files were carried out using Spats v2.0.5 and run with Python 2.7 through Anaconda 2.4. Additional options used during Spats analysis are provided in Supplementary Information.

### Generating secondary structure predictions

Preliminary secondary structure predictions for informing SHAPE-Seq experiments were generated using RNAstructure version 6.3 from the Mathews Lab (https://rna.urmc.rochester.edu/RNAstructure.html) (49).

SHAPE-Seq-informed secondary structure predictions were generated using Reconstructing RNA Dynamics from Data (R2D2) with the protocol and code detailed in methods of Yu, et al. (42). Additional options used during R2D2 analysis are provided in Supplementary Information. Predicted structures were visualized as bowplots using RNAbow software (50) and secondary structures using VARNA version 3.9 (http://varna.lri.fr/) (51).

### Template-based homology and molecular dynamics modeling of the *yxjA* riboswitch

To model the aptamer states, recently solved X-Ray Free Electron Laser (XFEL) structures of the *pbuE* adenine riboswitch aptamer (PDB IDs 5E54 and 5SWE) were used as templates to model the apo1, apo2 and holo states of the *yxjA* aptamer. The sequence of the *yxjA* aptamer was obtained from the *B. subtilis* genomic sequence. All modeling was performed using the Molecular Operating Environment (MOE) software package (52). In MOE, the *yxjA* aptamer sequence was aligned with the *pbuE* aptamer sequence such that the length of the conserved regions was identical (nucleotides 8 – 50 and 53 – 74), and such that the two additional nucleotides 51 – 52 in the *yxjA* aptamer fell in the variable L3 loop region. The *pbuE* aptamer sequence was then mutated in place based on secondary structure predictions generated by R2D2. For the apo1 state, chain A of PDB structure 5E54 was used. For the apo2 state, chain B of PDB structure 5E54 was used. For the holo state, chain A of PDB structure 5SWE was used.

To model the ligand binding pocket, a guanine nucleobase was manually inserted into the 3WJ of the modeled *yxjA* holo aptamer structure in MOE such that all proposed hydrogen bonding and stacking interactions were represented. A previous study showing that the global structure of the guanine binding pocket is nearly identical to that of the adenine binding pocket was used as a guide (53). Bound guanine is held in place with the same network of hydrogen-bonding interactions, and C74 is the specificity-determining pyrimidine. Additionally, there are two conserved recognition triples that stack on top of each other to organize the aptamer in response to bound guanine.

To model the expression platform, sequence and secondary structure information for nucleotides 1 – 7 and 75 – 124 that contain the remainder of the P1 helix, a section of the P4 helix, and the rest of the nucleotides that form the central helix was obtained from secondary structures of the full length terminated and antiterminated conformations, respectively. This information was provided as input into the open-source tertiary structure generator RNAComposer (54). We narrowed down the 60 conformations predicted by RNAComposer based on two criteria: (1) whether the base pairs involved in central helix formation were equidistant between their P1 and central helix complements and (2) whether the conformation showed good agreement with the R2D2-based secondary structure predictions of the 160 nt intermediate state. The existing apo2 and holo aptamers was stitched onto the chosen 53 nt expression platform conformation in MOE to construct 124 nt hybrid models. Additionally, we created a separate holo hybrid model with a U80G mutation to mimic the broken ‘ON’ mutant (U91G in Figure 4C) that was studied experimentally.

### All-atom molecular dynamics simulations

All-atom molecular dynamics (MD) simulations were performed using the GROMACS 2019.4 software package (55); RNA was represented by the Amber-99 force field (56) with Chen-Garcia modifications for RNA bases (57) and Steinbrecher-Case modifications for the backbone phosphate (58). Each structure was placed in the center of a cubic box such that there was at least 1.5 nm from all RNA atoms to the box edge in each direction, then solvated with sufficient TIP4P-EW water and K^+^ and Cl^-^ ions to mimic 1 M excess salt conditions. The systems were then energy minimized using the steepest descent algorithm for 10,000 steps with a 2 fs timestep and a force tolerance of 800 kJ mol^-1^nm^-1^. Once energy minimized, all systems were simulated in the *N,P,T*-ensemble using a leapfrog Verlet integrator with a time step of 2 fs. An isotropic Berendsen barostat was used for pressure coupling (59) at 1 bar with a coupling coefficient of τ_p_ = 1 ps, and a velocity-rescale Berendsen thermostat was used for temperature coupling (60) with a time constant of τ_t_ = 0.1 ps. The LINCS algorithm was used to constrain all bonds, and a cutoff of 1.0 nm was employed using the Verlet scheme for Lennard-Jones and Coulomb interactions.

The apo1, apo2 and holo aptamer models were equilibrated at 310 K for a total of 50 ns. Base pairs predicted using R2D2 were enforced by applying distance-dependent piecewise flat-bottomed harmonic bias restraints between the central hydrogen bond donor and acceptor of paired bases at a strength of 5 kcal/mol. After equilibration, all aptamer models were simulated at a wide range of temperatures (310 K – 400 K) for a total of 50 ns without any restraints.

The 124 nt apo2 and holo hybrid models were also equilibrated at 310 K for a total of 50 ns. Base pairs in the AD predicted using R2D2-generated secondary structure predictions were reinforced by applying harmonic bias restraints at a strength of 5 kcal/mol.

### Two-dimensional replica exchange molecular dynamics simulations

After equilibration, two-dimensional replica exchange molecular dynamics (2D REMD) simulations were run on the apo2 and holo hybrid models to directly probe the strand exchange process. The first series of 2D REMD simulations were performed using nine replicas at temperatures 373 K, 374 K, and 375 K. Harmonic bias restraints were applied at strengths of 4.5 kcal/mol to base pairs 75A – 107U, 76A – 106U, and 77A – 105U, and 5 kcal/mol to base pairs 78G – 104C, 79U – 103A, and 81U – 101A. As a result of this biasing, successful strand invasion was observed under 50 ns frequently in the apo2 hybrid model but infrequently in the holo hybrid model (Supplementary Table S6). As such, these initial simulations were run for 22 ns each, for a total of 198 ns per simulation. To further confirm what was observed in these simulations, a second series of 2D REMD simulations was performed using nine replicas at temperatures 373 K, 374 K, and 375 K. Harmonic bias restraints were applied at strengths of 4 kcal/mol to base pair 74G – 108, 4.5 kcal/mol to base pairs 75A – 107U, 76A – 106U, and 77A – 105U, and 5 kcal/mol to base pairs 78G – 104C, 79U – 103A, and 81U – 101A. Again, due to this biasing, central helix formation was observed under 50 ns frequently in the apo2 hybrid model but infrequently in the holo hybrid model (Supplementary Table S6). These simulations were run for 47 ns each, for a total of 423 ns per simulation. Finally, a third 47 ns 2D REMD simulation was performed using nine replicas with identical temperatures and the same gradient of harmonic bias restraints to probe the strand exchange process in our broken ON (U80G in hybrid model, U91G in *in vivo* assays) holo hybrid model. Altogether, the cumulative 2D REMD simulation time amounted to ∼1.6 µs.

## RESULTS

### The *yxjA* riboswitch reporter construct downregulates GFP expression in response to ligand

We began our studies by developing a cellular assay to evaluate the function of the *yxjA* riboswitch. We assembled several versions of an expression construct consisting of the *Escherichia coli* sigma 70 consensus promoter, a variable portion of the pre-aptamer leader sequence, an adenine-responsive *Bacillus subtilis yxjA* riboswitch sequence, and the superfolder GFP (sfGFP) coding sequence (Supplementary Figure S1). This construct contained a mutation at position 79 to make the riboswitch responsive to adenine and adenine analogs (G79T labeled as WT) based on prior studies that demonstrated guanine riboswitches with this mutation have a high binding affinity for 2-aminopurine (2AP) (43). In addition to the wildtype aptamer sequence, we also designed a ligand-unresponsive *yxjA* mutant (C79A) to serve as a control. Constructs were transformed into *E. coli* TG1 electrocompetent cells, grown overnight in LB media, then used to inoculate subcultures in M9 minimal media with or without 2-aminopurine (2AP) ligand added.

We first optimized subculture growth time by collecting a time course of SFGFP fluorescence data over six hours of subculture growth to identify the optimal time point that gave the maximum difference in fluorescence between the with and without ligand conditions. We found this time point to lie at five hours after subculture inoculation (Supplementary Figure S2). After determining this optimal time point, we collected cellular growth data at the same time point to test for potential ligand toxicity. This experiment determined that extracellular concentrations of 1 mM 2AP in the media caused some changes in cellular growth compared to the 0 mM 2AP condition but was still tolerable for our assay conditions (Supplementary Figure S3). An additional point of construct optimization involved examining the 5′ pre-aptamer leader sequence that is encoded before the aptamer domain (AD). Initial studies of this construct architecture demonstrated that introducing an additional five nucleotides of the pre-aptamer leader sequence provided the maximal dynamic range of ligand response over other lengths (Supplementary Figure S4). Finally, we used this optimized construct and assay conditions to characterize the dose response of the riboswitch construct to a range of ligand concentrations (Figure 1B). Collectively, these results demonstrate that our reporter system and assay conditions can characterize the function of the *yxjA* riboswitch in *E. coli* cells.

### The P1 helix of the *yxjA* riboswitch has specific length requirements for ligand-responsive activity

Once our cellular function assay was established, we next investigated the dependence of *yxjA* riboswitch function on the length of the aptamer P1 helix. Previous work has shown across multiple riboswitches that the P1 helix is a critical site for controlling the ligand-responsiveness of transcriptional riboswitches (61). This is particularly true for riboswitches such as the *pbuE* adenine or tetrahydrofolate (THF) riboswitches where ligand binding occurs within junctions, and the P1 stem not only helps to maintain the base of the ligand binding pocket, but also shares direct overlapping sequence with the EP (62). In these cases, this overlapping sequence maintains mutual exclusivity between the terminated and anti-terminated structures. However, at the secondary structure level in the *yxjA* riboswitch, the terminator can fully form when the P1 helix is only partially disrupted, leaving the AD and ligand binding pocket intact (Figure 1C and D). We therefore wanted to investigate if P1 length is important in this architecture, and if it would reveal details about the switching mechanism. To test this idea, we designed mutants that altered the 5′ side of P1 to disrupt base pairing while preserving base pairs involved in terminator formation.

Characterization of a range of P1 shortening and lengthening mutations showed that the optimal P1 length for the *yxjA* riboswitch to function lies between 7 (WT-6) and 14 (WT+1) base pairs. As P1 base pairs are removed, mutant riboswitches functioned similarly to the WT until seven base pairs remained (Figure 2A). At six base pairs or shorter, SFGFP fluorescence remained relatively high even in the presence of ligand, representing a riboswitch broken in the ‘ON’, or anti-terminated structure. As P1 base pairs were lengthened, the mutants functioned similarly to WT only up until one additional base pair (Figure 2B). Any P1 helix longer than 14 base pairs was characterized by low SFGFP expression even without ligand present, which we interpreted to mean that the riboswitch was broken in the ‘OFF’ or always terminated structure.

**Figure 2.**
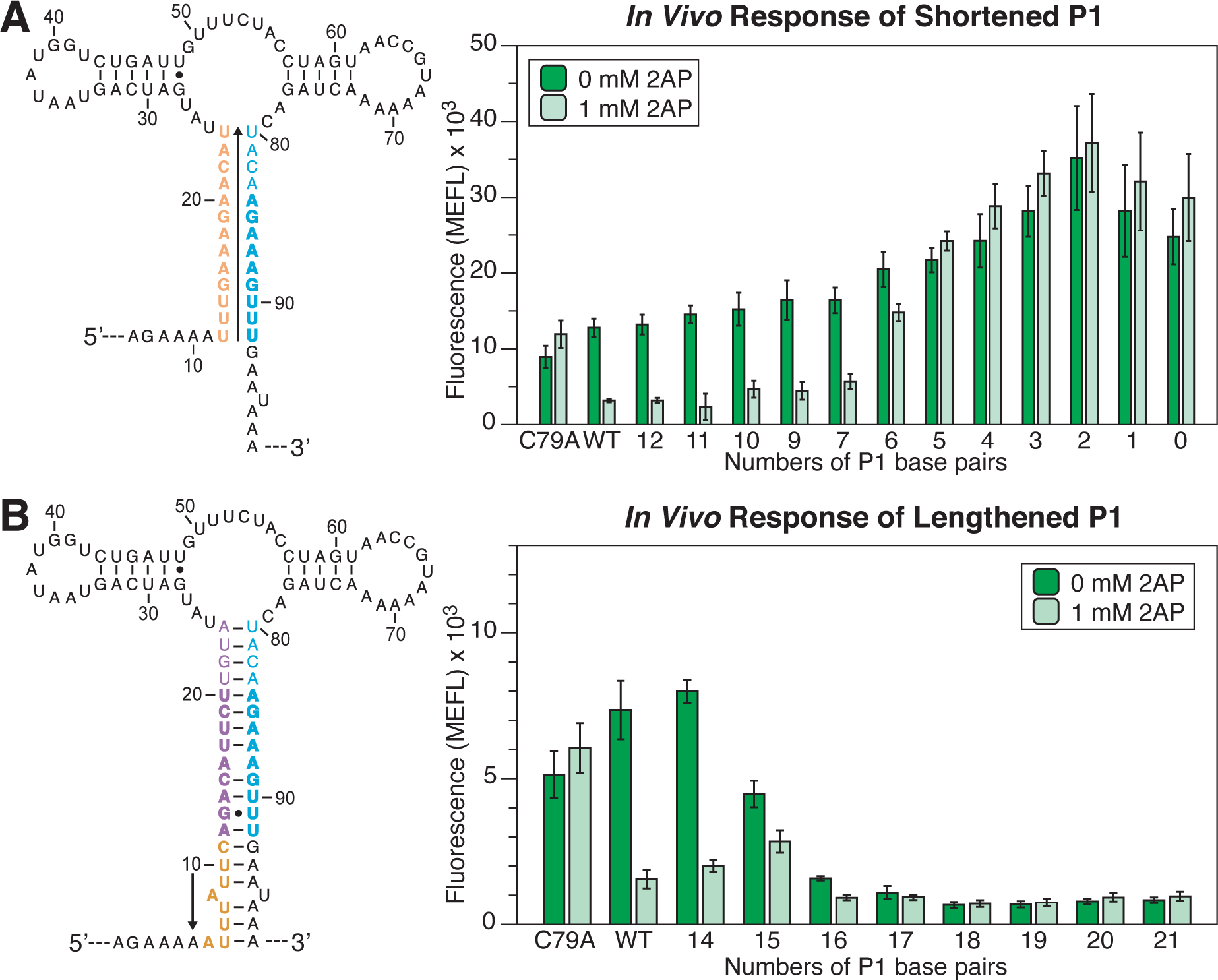
Flow cytometry gene expression characterization data of P1 mutants. **(A)** Shortened P1 mutants were designed by mutating the 5′ side of the P1 helix into the same nucleotide identity as the 3′ side to prevent base pair formation (orange). Mutations were introduced in successive order beginning from the bottom of P1 towards the 3WJ. **(B)** Lengthened P1 mutants were designed by successively inserting additional nucleotides on the 5′ side of the P1 (orange) upstream of the riboswitch AD and downstream of the pre-aptamer sequence, to lengthen the P1 stem. Data was collected as described in Figure 1 using the indicated amount of ligand (2AP) in the media. Bars represent averages from three biological replicates, each performed in triplicate technical replicates for a total of nine data points (n=9), with error bars representing standard deviation.

These results suggest that the P1 helix region between 7 and 14 base pairs plays an important role in the switching mechanism. Since this region shares sequence overlap with the terminator (Figure 1C and D), we hypothesized it could play an analogous role to the “overlap region” between the AD and EP that ensures exclusivity between the anti-terminated and terminated structures in other riboswitches. This overlap region differs from those in the *pbuE* riboswitch and other ‘ON’ riboswitches that extend into the direct ligand binding portion of the AD. Interestingly, for *yxjA* to function, it requires P1 to be at least seven base pairs long, which is longer than the minimum three base pairs required for function in the *pbuE* riboswitch (63). Overall, this suggests that the overlap region is critical for facilitating the *yxjA* regulatory mechanism.

### Modeled structures based on chemical probing data suggest a possible mechanism involving strand exchange

To investigate the structural intermediates that mediate the *yxjA* riboswitch regulatory mechanism, we performed cotranscriptional *in vitro* Selective 2’-Hydroxyl Acylation Analyzed by Primer Extension Sequencing (cotranscriptional SHAPE-Seq) experiments using benzoyl cyanide (BzCN) to chemically modify the RNA at more flexible residues during stalled transcription at various lengths (27, 64). The resulting chemical probing reactivity data was then used as inputs into the Reconstructing RNA Dynamics from Data (R2D2) algorithm to generate experimentally informed models of cotranscriptionally folded structural intermediates (27, 42). Our goal with these experiments was to observe evidence of a central helix – a structure formed by the complementary regions of P1 and the terminator (Figure 1C) – that could form during the folding pathway, or an alternative structural model that we could use to further investigate the riboswitch mechanism. As such, we focused these experiments on transcription conditions without ligand.

We selected three distinct lengths of the riboswitch: before where strand invasion by the central helix should occur, after where strand invasion by the central helix should have occurred, and the full length riboswitch. We generated secondary structure predictions using RNAStructure (49) to help inform our selection (Supplementary Figure S6) and designed purified DNA templates that would allow the *E. coli* RNA Polymerase (*Ec*RNAP) to transcribe the riboswitch up to a length of 135 nt (before the central helix has invaded P1), 160 nt (central helix has invaded part of P1), and 220 nt (the full length of the riboswitch).

We chemically probed these lengths of the *yxjA* riboswitch in stalled transcription elongation complexes with *E. coli* RNA polymerase (*Ec*RNAP) and tested two replicates for each length. For each transcribed RNA, we included an additional 14 nt in the DNA template design to account for the *Ec*RNAP footprint (65). We also performed additional equilibrium-refolding experiments where the RNA was synthesized, purified, denatured, and equilibrium refolded in the absence of *Ec*RNAP, and therefore did not include the additional 14 nt. Reverse transcription and high-throughput sequencing allowed us to uncover SHAPE reactivity information for each nucleotide of these transcripts. Our resulting SHAPE data revealed general consistencies in reactivity patterns between replicates (Supplementary Figure S7 and S8). We then used the SHAPE data to experimentally inform computational structure models with R2D2 to uncover details of cotranscriptional structures that may not be detectable by minimum free energy (MFE) structure predictions (42).

For each RNA length, R2D2 generated 100 structures that are most consistent with the observed chemical probing data. These data are conveniently viewed as RNAbow plots (50) that capture a population of RNA structures. In these diagrams, arcs between two positions indicate a base pair between those two positions and the opacity of the arc line indicates how frequent that pair occurred in the population of structures. Viewing the RNAbow plots from our cotranscriptional SHAPE-Seq experiments of these lengths revealed the presence of the P1 helix at length 135 nt, the potential presence of a central helix at length 160 nt, and a possible terminated structure at full length (Figure 3).

**Figure 3.**
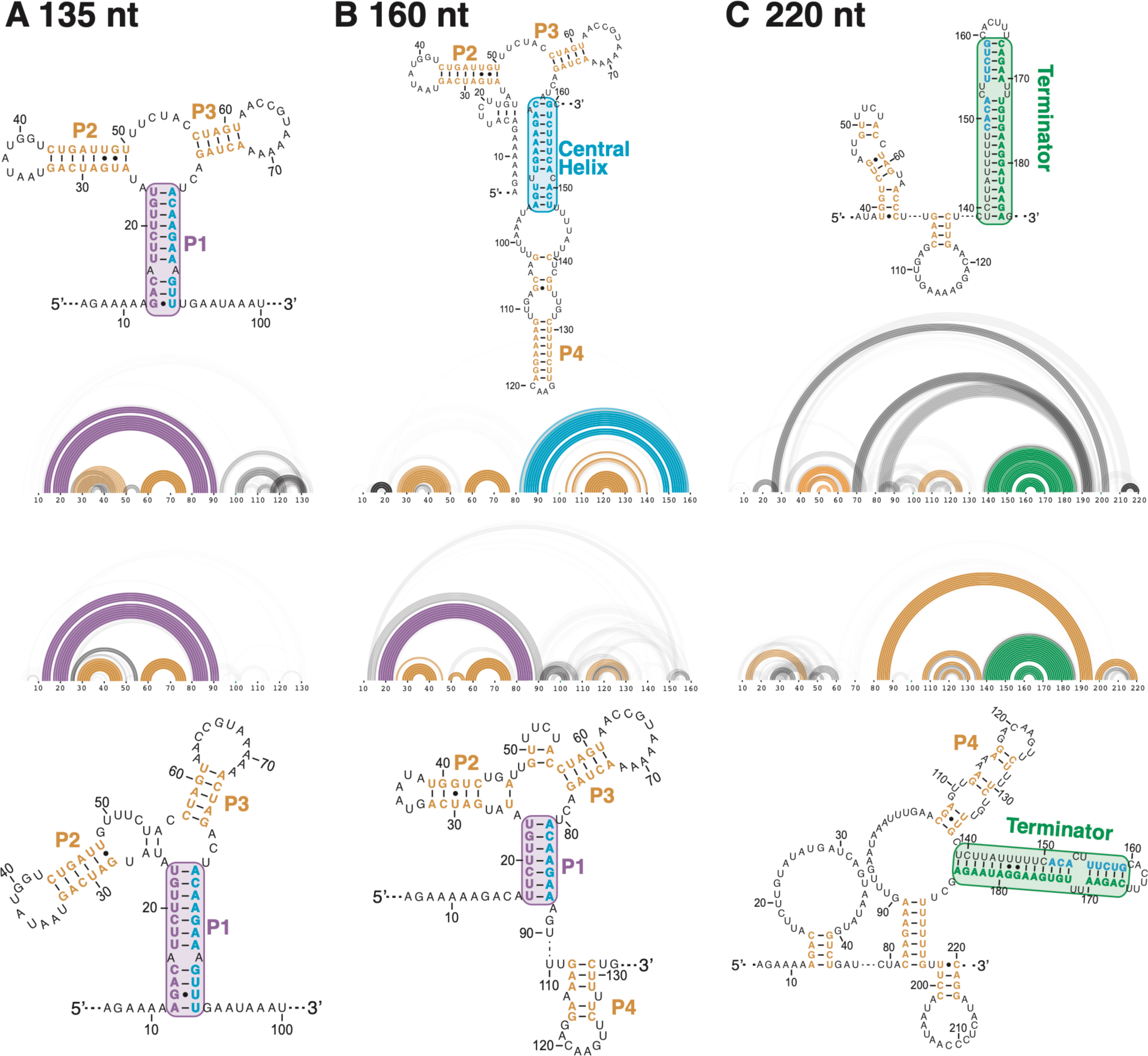
Experimentally informed models of structural intermediates along the *yxjA* folding pathway without ligand. Cotranscriptional SHAPE-Seq experiments were performed at two intermediate lengths of **(A)** 135 nt, **(B)** 160 nt, and **(C)** for the full length riboswitch (220 nt) in the absence of 2AP. Reactivity data were then incorporated into R2D2 pathway modeling to generate 100 structures that are maximally consistent with the reactivity data. This family of structures is displayed as RNAbow plots, where an arc between two positions indicates a base pair in a specific structure, and the opacity of the arc indicates the prevalence of the base pair among the selected structure. Two replicates were performed at each length. The consensus structure over the population of selected structures is shown as a secondary structure alongside each bow plot. In the secondary structures, the central helix is colored teal while the 5′ side of the P1 helix is purple and the 3′ side of the terminator is green. Colored boxes drawn around the P1 helix, the central helix, or the terminator, as well as the color of the P2/P3/P4 helices, match the color of arcs in the RNAbow plots.

**Figure 4.**
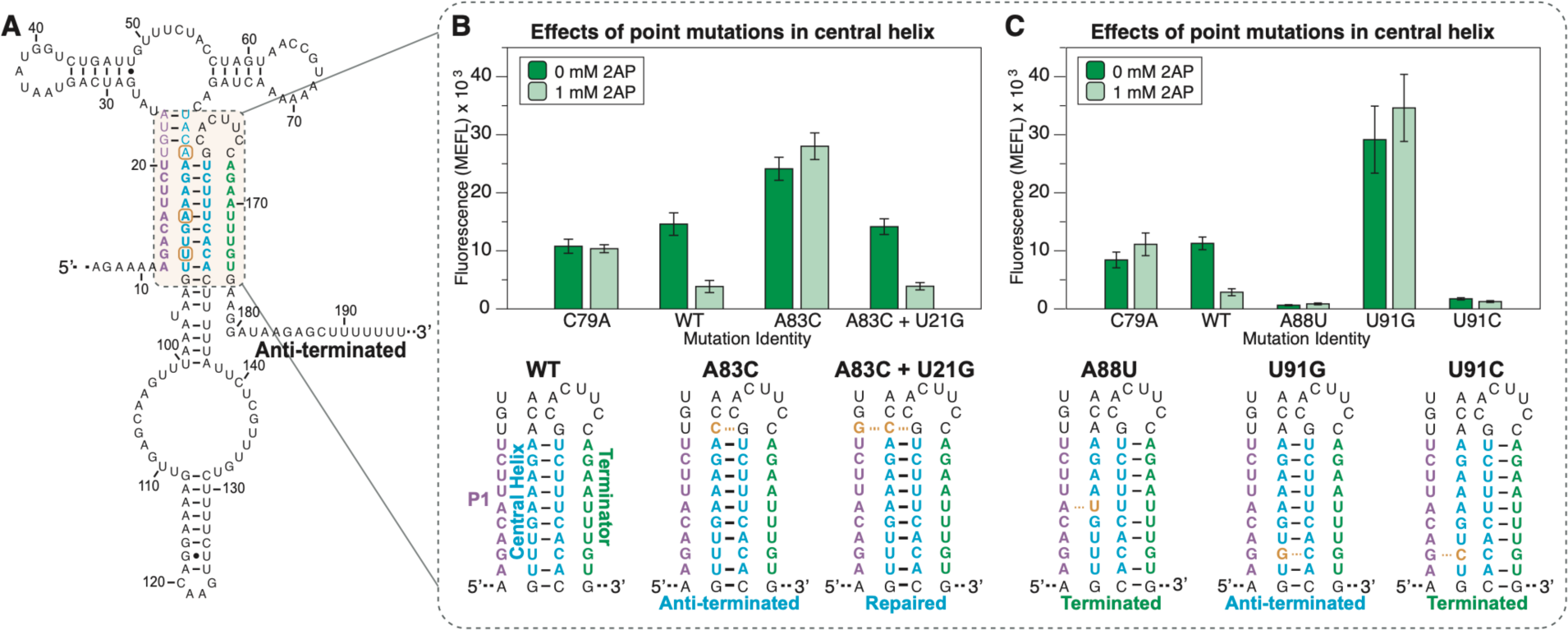
Flow cytometry gene expression characterization data of *yxjA* riboswitches containing point mutations and rescue mutations in the central helix. **(A)** Schematic of mutation locations (orange circles) based on where natural single mismatches were found. **(B)** Single mismatch mutants at position 83 (A83C) and repair at position 21 (A83C + U21G), and **(C)** single mismatch mutants at positions 88 and 91. Schematic of mutations are shown below the bar graphs. Data was collected as described in Figure 1 using the indicated amount of ligand (2AP) in the media. Bars represent averages from three biological replicates, each performed in triplicate technical replicates for a total of nine data points (n=9), with error bars representing standard deviation.

At length 135 nt, we observed that most structures in the population indicated P1 formation, as well as base pairs in P2 and P3 (Figure 3A). We saw similar features in the equilibrium data, with the additional presence of the P4 stem (Supplementary Figure S9A). At length 160 nt, we saw differences between our two replicates. One replicate showed the formation of P2, P3, and a central helix, resembling an anti-terminated structure. The second replicate still contained an intact P1, along with P2 and P3 in the AD and the P4 stem (Figure 3B). In the equilibrium data, both replicates revealed the presence of the central helix (Supplementary Figure S9B). Finally, at the full length, we saw the base pairs of the previously predicted terminator structure reflected in the R2D2 output, but also the presence of other sets of base pairs that did not correspond to previously predicted or proposed secondary structure features (Figure 3C). The equilibrium data contained similar features, most notably the terminator (Supplementary Figure S9C). We suspected a lower read depth in the full lengths due to experimental conditions may have made these results less clear for interpretation.

Overall, these results suggest that an intermediate structure consisting of a central helix that competes with part of the P1 helix sequence can form during transcription. The presence of the terminator at full length may also imply that the terminator may need to invade through the central helix, or compete with it, to fully form as the final structure.

Based on this structural data, we therefore hypothesized that the *yxjA* riboswitch mechanism depends on a competition of two strand exchange processes to regulate gene expression: formation of the central helix via strand displacement of the P1 stem, and formation of the terminator via strand displacement of the central helix. Depending on which strand exchange process can properly form base pairs (central helix vs. terminator), the more favored helix will determine which of the two mutually exclusive final structures the riboswitch will adopt.

### Naturally encoded mismatches in the central helix control riboswitch response to ligand binding

We next sought to investigate how mutations in key regions may affect strand invasion in our hypothesized riboswitch mechanism. The region of the P1 helix shown to be critical for function in our previous length requirement experiments (Figure 2) encodes single mismatches that are not expected to form base pairs in either the P1 or central helix in secondary structure predictions (Figure 4A). Mismatches within helical regions are predicted to slow the progression of strand displacement mechanisms by introducing a barrier that stalls the progress of an “invading” strand from gaining an additional base pair with the “substrate” strand and displacing the “incumbent” strand (66). Depending on where they occur, these mismatches may slow the formation of the P1 helix or the central helix and bias the folding pathway accordingly. We therefore hypothesized that if we repaired a mismatch to instead form a base pair within the P1 helix, it would favor P1 formation and outcompete central helix formation to facilitate terminator formation and fold into a broken ‘OFF’ state. Correspondingly, if we mutated the mismatch to form a base pair in the central helix instead, favoring the central helix would inhibit terminator formation and lead to an always ‘ON’ phenotype.

We tested this hypothesis by making several point mutations and characterizing riboswitch function. We found that when we mutated a single nucleotide to form a base pair with either P1 or the central helix, the riboswitch would either break ‘ON’ or ‘OFF’ according to our hypothesis, regardless of ligand in the media. For example, extending the central helix by a single base pair by mutating position 83 from an A to a C broke the riboswitch into the ‘ON’ phenotype (Figure 4B). When the opposing nucleotide at position 21 was mutated to a G so that both P1 and the central helix could form this pair, the riboswitch function was restored (Figure 4B).

In another example, positions 88 and 91 encode an A:A mismatch or G:U wobble pair, respectively, in the P1 helix. In this case, the A88U mutation, which allows a base pair to form within P1, but makes the corresponding pair in the central helix a mismatch, broke the riboswitch into the ‘OFF’ phenotype (Figure 4C). A similar pattern held true for position 91 which can normally form a wobble pair within P1. When mutated to cause a G:G mismatch in P1, the results yield a broken ‘ON’ phenotype. But when mutated to cause a strong G:C base pair in P1 and a broken C:C pair in the central helix, the alteration yields a broken ‘OFF’ riboswitch (Figure 4C).

These results demonstrate that these naturally occurring P1 and central helix mismatches are critical for determining the regulatory outcomes of the *yxjA* riboswitch. By altering them to form base pairs in either the P1 or the central helix, we could distinctly favor the P1-terminator structure or the central helix/anti-terminator structure. This observation also suggests that the ability to strand exchange plays an important role in the riboswitch folding process and that mismatches may introduce a stalling effect on invading strands.

### Central helix base pairing is required for riboswitch function

After discovering specific requirements for the P1 helix length and single mismatches that could completely shift riboswitch function in one direction or the other, we next designed mutants to test the importance of base pairing in the central helix structure for riboswitch function. We created these mutants by introducing broken base pairs in the central helix to observe how breaking multiple continuous base pairs would affect riboswitch function. Based on observations from our previous results, we hypothesized that breaking these base pairs would prevent the central helix from forming and disrupt riboswitch ligand response, favoring the formation of the terminator and leading to a broken ‘OFF’ phenotype. To test this hypothesis, we introduced these mutants by swapping the base pairs in the terminator so the central helix was broken while still maintaining base pairing in the terminator (Figure 5).

**Figure 5.**
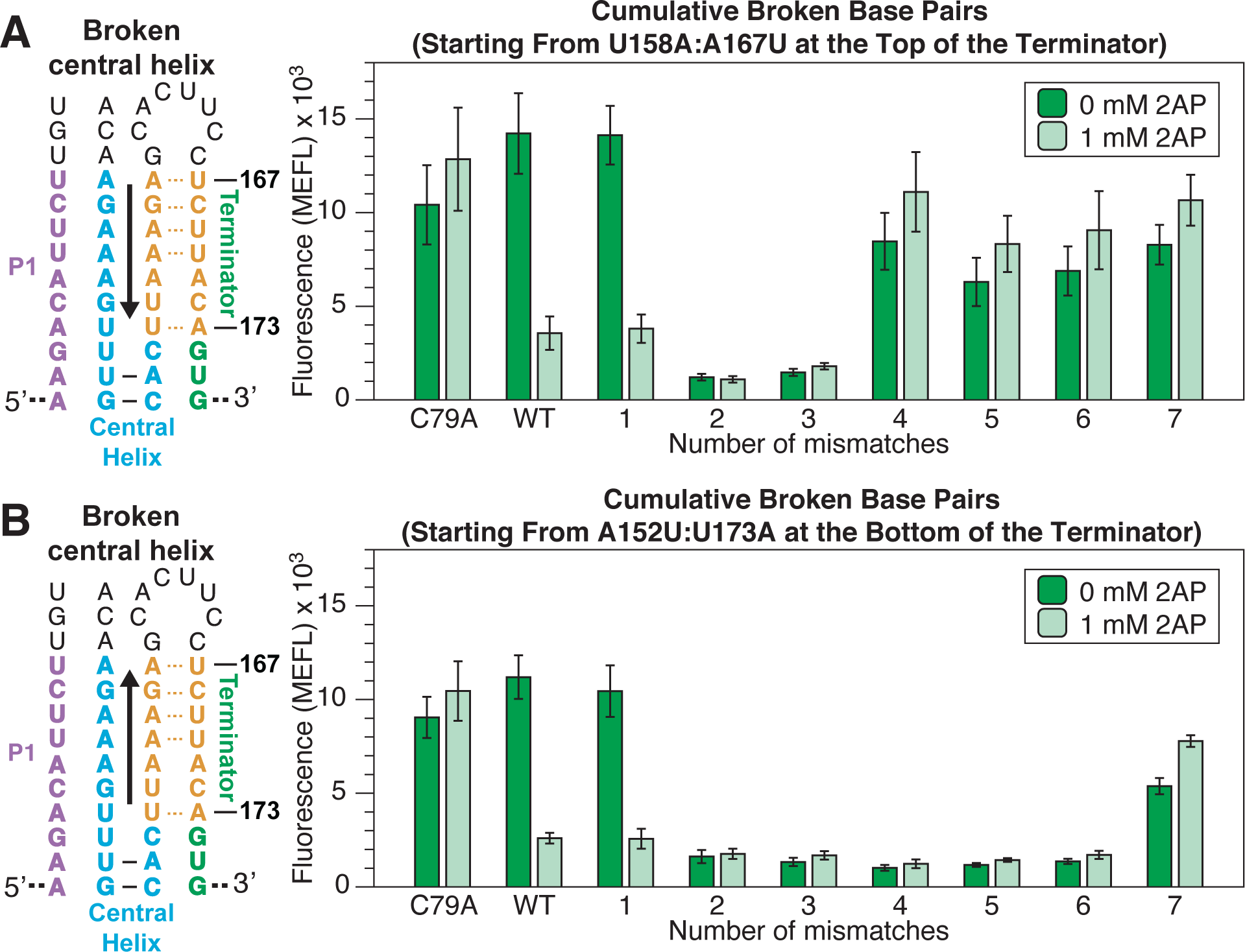
Flow cytometry gene expression characterization data of *yxjA* riboswitches containing mismatches along the central helix. **(A)** Mismatches (orange) were designed to interrupt sequential base pairs from the top of the central helix by swapping the 5′ and 3′ side of the terminator to break the base pair in the central helix while maintaining the pair in the terminator. The number of sequential mismatches is indicated under the characterization data. **(B)** Mismatches designed to interrupt sequential base pairs from the bottom of the central helix were designed in a similar manner. Data was collected as described in Figure 1 using the indicated amount of ligand (2AP) in the media. Bars represent averages from three biological replicates, each performed in triplicate technical replicates for a total of nine data points (n=9), with error bars representing standard deviation.

As hypothesized, when we introduced these broken base pairs, the riboswitch displayed a broken ‘OFF’ phenotype. When the broken base pairs were introduced starting from the top of the central helix (U158A:A167U), the riboswitch could only tolerate one broken base pair before breaking ‘OFF’ even in the presence of ligand (Figure 5A). Introducing broken base pairs from the bottom (A152U:U173A) yielded the same results (Figure 5B).

However, we noticed that introducing more base pair swaps in the terminator caused the riboswitch to display a broken ‘ON’ phenotype. We hypothesized that this was due to introducing too many sequence changes that ultimately impaired terminator function. To test this hypothesis, we designed constructs that removed the AD to test the terminator’s baseline ability to terminate without interference from a competing AD. This experiment allowed us to test the function of mutated terminators with the same cellular gene expression assay. As expected, the terminator experiments showed that termination was impaired when either four or more swaps were introduced from the top (U158A:A167U) (Supplementary Figure S10), or seven swaps were introduced from the bottom (A152U:U173A), matching the same regions that created a broken ‘ON’ phenotype in the swapped riboswitch constructs (Supplementary Figure S11).

Overall, these results demonstrate that base pairing within the central helix is critical for riboswitch function.

### All-atom molecular dynamics simulations suggest the P1 helix becomes more conformationally rigid upon ligand binding

Our data suggests that the *yxjA* regulatory mechanism is mediated by the competition between the formation of the P1 helix, and the central helix. A full mechanistic model of the regulatory mechanism would then need to explain how ligand binding biases the initial folding of the P1 helix structure towards the ‘OFF’ functional state. Based on our mutational results, we hypothesized that ligand binding to the AD junction prevents the central helix from invading a fully formed P1, though our functional data is not able to explain the structural basis for P1 stabilization. To investigate this further, we conducted all-atom molecular dynamics (MD) simulations to probe specific structural differences between the apo and holo states of the *yxjA* aptamer.

Prior time-resolved crystallographic studies investigated adenine-binding to the *pbuE* riboswitch aptamer and found the 3WJ and P1 helix differs between the apo and holo aptamer states (67). Specifically, this study showed that in the holo form, the P1 helix has less conformational freedom compared to that observed in the apo form. Due to the high level of sequence structural conservation between adenine and guanine riboswitches (45-47), we hypothesized that the same difference in P1 helix flexibility observed in the apo and holo states of the *pbuE* aptamer would be observable in the apo and holo states of the *yxjA* aptamer. To test this hypothesis, we used the X-ray crystal structures of the two distinct *pbuE* aptamers in apo conformation as templates to construct homology-based models for two *yxjA* apo conformations: apo1 and apo2 (67). We also used the *pbuE* crystal structure to construct the *yxjA* holo aptamer state. We used these three models as our starting points for all-atom MD simulations for 50 ns at 310 K. We aligned the resulting trajectories from each simulation, using the initial structure of the AD (aside from the P1 helix) as the reference state. To analyze these aligned trajectories, we overlaid every 500^th^ frame of the 3′-strand of the P1 helix, which is involved in base pair formation in both the terminated and anti-terminated conformations (Figure 6A). We also calculated the per-residue root mean square fluctuation (RMSF) of the 3′-strand of the P1 helix to quantify the average flexibility of this region over time (Table 1).

**Figure 6.**
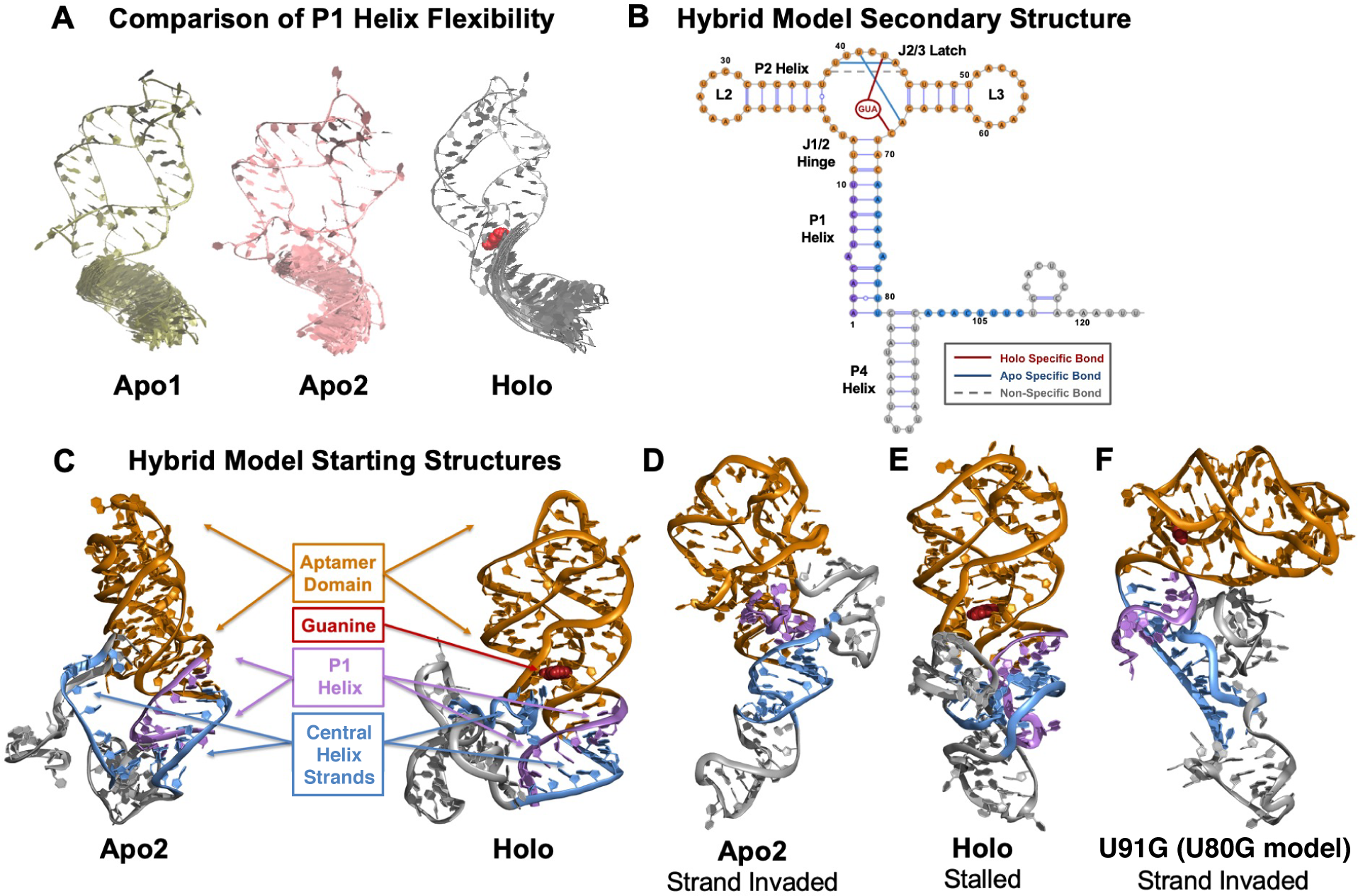
Snapshots of MD and 2D REMD simulations. **(A)** A visual comparison of every 500^th^ frame of the 3’-strand of the P1 helix in the apo1, apo2 and holo aptamers after a 50 ns MD simulation shows increased rigidity from the apo states to the holo state. **(B)** The secondary structure of the hybrid model of the *yxjA* riboswitch used in MD simulations. Hydrogen bonds that form in apo and holo states within the aptamer junction region are annotated, alongside interactions with the ligand. **(C)** Three-dimensional starting structures of the 124 nt hybrid model of the apo2 and holo states after a 50 ns equilibration. Different domains are color coded as indicated. **(D)** An ending conformation of one 2D REMD simulation of the apo2 hybrid model showing successful strand exchange of the P1 helix to form the central helix. **(E)** An ending conformation of one 2D REMD simulation of the holo hybrid model showing a stalled strand exchange process that leaves the P1 helix intact. **(F)** An ending conformation of one 2D REMD simulation on the broken ‘ON’ U91G mutant (U80G in hybrid model) holo hybrid model showing successful strand exchange of the P1 helix to form the central helix.

**Table 1.** Per-residue root mean square fluctuation (RMSF) of the 3′-strand of the P1 helix in the apo1, apo2 and holo aptamer states after a 50 ns MD simulation.

| **P1 Helix Residue** | **Apo1**<br>**RMSF (Å)** | **Apo2**<br>**RMSF (Å)** | **Holo**<br>**RMSF (Å)** |
| --- | --- | --- | --- |
| 69 | 1.380 | 1.468 | 1.246 |
| 70 | 1.041 | 1.022 | 0.807 |
| 71 | 1.002 | 0.819 | 0.695 |
| 72 | 1.035 | 0.819 | 0.708 |
| 73 | 0.959 | 0.847 | 0.687 |
| 74 | 0.914 | 0.937 | 0.696 |
| 75 | 1.019 | 0.995 | 0.814 |
| 76 | 1.446 | 1.337 | 0.121 |

These simulation results show a clear difference in P1 helix flexibility between apo and holo states. Specifically, in the apo1 state, we observed a range of simulation conformations of the P1 helix and the largest RMSF distances, indicating that the P1 helix is flexible and dynamic in this state. In contrast, in the holo state, there is a distinct reduction in possible P1 conformations with reduced RMSF distances of approximately 0.2 – 0.3 Å in the RMSF of residues 70 – 74 in the holo state. We found that the apo2 state is in between the apo1 and holo states, with slightly reduced conformational flexibility from the apo1 state, indicating that the helix is more compressed. From these results, we can conclude that, in agreement with the observations for the *pbuE* aptamer, the binding of guanine to the *yxjA* aptamer causes the P1 helix to become more rigid, particularly in the 3′-strand of P1, which is involved in the formation of the central helix during strand exchange.

### All-atom MD simulations of hybrid models of the *yxjA* riboswitch suggest that a more conformationally rigid P1 helix interrupts strand exchange by the central helix

Having established that ligand binding can alter the flexibility of the P1 stem, we next used two-dimensional replica exchange molecular dynamics (2D REMD) simulations to investigate the strand exchange process by which the central helix is formed in place of the P1 helix in the ligand-free apo state. To directly probe the process of strand exchange, we designed a 124 nt model of the *yxjA* riboswitch, consisting of both the AD and an intermediate state of the EP. The remainder of the P1 helix was taken from a secondary structure prediction of the 195 nt terminated conformation, while the P4 helix was adapted from a secondary structure prediction of a 195 nt antiterminated conformation (Figure 6B). To generate the three-dimensional structure of the EP, we provided the sequence and secondary structure information for our 124 nt intermediate as input into the open-source tertiary structure generator RNAComposer (54). We narrowed down the 60 conformations predicted by RNAComposer based on two criteria: (1) whether the base pairs involved in central helix formation were equidistant between their P1 and central helix complements and (2) whether the conformation showed good agreement with the R2D2-based secondary structure predictions of the 160 nt intermediate state. We then stitched our existing apo2 and holo aptamers onto our chosen EP conformation using Molecular Operating Environment (MOE) (52). Finally, we ran single-temperature all-atom MD simulations on our final 124 nt apo2 and holo hybrid structures at 310 K for a total of 50 ns to equilibrate the AD and the expression platform (Figure 6C). We enforced base pairs at the start of each helix, at the end of each helix, the kissing loop interactions, and in the binding pocket of the AD by applying distance-dependent harmonic bias restraints (Supplementary Figure S12).

With the model established, we next aimed to identify the mechanisms by which bound guanine in the 3WJ rigidifies the P1 helix. We first ran all-atom MD simulations of our equilibrated apo1, apo2, and holo aptamers at a wide range of temperatures (310 K – 400 K) for 50 ns each (Supplementary Video S1). Again, we used per-residue RMSF to quantify average flexibility over time versus temperature. Our RMSF analysis revealed that the J1/2 hinge, which is thought to serve as a pivot point for the P1 helix (67), is nearly three times more flexible in apo1 and apo2 as compared to holo (Supplementary Figure S13A). This observation suggests that bound guanine to the 3WJ tethers the J1/2 hinge to the AD such that it can no longer serve as a point of rotation for the P1 helix. Similarly, the J2/3 latch is found to be nearly twice as flexible in apo1 and apo2 as compared to the holo state due to the formation of base triplets and stacking interactions that occur in the binding pocket upon ligand recognition (Supplementary Figure S13B).

Following the same trend, the kissing loop interaction between L2 and L3 is significantly more stable in the holo form than either apo1 or apo2, where it is dynamic and constantly dissociating and reassociating (Supplementary Figure 13C). As the only difference between the two simulations is the presence of bound guanine (i.e. no restraints were placed on the kissing loop interaction for either state), this effect must be a direct result of increased binding pocket organization upon recognition of guanine. In contrast to the overall trend, we found that the P3 helix shows greater flexibility in the holo state than it does in the apo states, which is most noticeable at the P3-L3 interface, where some residues show a three-fold increase in RMSF (Supplementary Figure S13D). This increase is a direct result of the stabilized kissing loop interaction in the holo state. Conversely, the P2 helix exhibits greater stability in the holo state and increased flexibility in the apo1 and apo2 states (Supplementary Figure S13E).

Taken together, these results are in broad agreement with single-molecule pulling studies of the *xpt* guanine riboswitch (68), where increased kissing loop stability in the holo state led to increased flexibility in the P3 helix, especially at the P3-L3 interface, ultimately increasing P1 helix stability. The same study also observed the opposite trend for the apo state, where decreased kissing loop stability correlated with increased P3 helix stability and decreased P1 helix stability. We therefore conclude that this mechanism also holds true for the highly homologous *yxjA* riboswitch.

To observe the effect of bound guanine on the ability of the *yxjA* riboswitch central helix to undergo strand exchange, we ran a series of 2D REMD simulations on our 124 nt apo2 and holo hybrid models. To ensure that structurally plausible strand exchange events could be observed in feasible simulation timescales, we applied a gradient of harmonic bias restraints to the base pairs involved in the strand exchange process to form the central helix (Supplementary Figure S14). These restraints biased the model towards forming the anti-terminator conformation but were kept sufficiently weak that they would stall rather than unfold the AD if strand exchange was difficult to achieve. We hypothesized that the apo2 hybrid model would have a higher likelihood to successfully undergo strand exchange and form the anti-terminated conformation due to the increased flexibility of its P1 helix. Therefore, for the central helix to form, the 5′-end of the P1 helix would gradually dissociate from its starting position. Conversely, we predicted the holo hybrid model would not be able to form the anti-terminated conformation. Instead, the 5′- and 3′-ends of the central helix would move closer together to satisfy the distance restraints, but the structure would stall and would be unable to form the central helix. The starting P1 helix would remain largely intact, and strand exchange would not occur.

To test these hypotheses, we first ran a series of 2D REMD simulations (9 replicas, 22 ns per replica) on each hybrid model with a gradient of weak harmonic bias restraints applied to 6 central helix base pairs. The strongest harmonic bias restraint was applied to the lowermost base pair, and the weakest harmonic bias restraint was applied to the uppermost base pair (Supplementary Figure S14). Strand exchange was determined to be successful if at least 80% of the six restrained central helix base pairs formed over the course of a simulation. As hypothesized, we found that successful strand exchange occurred in the apo2 hybrid model on a consistent basis (Figure 6D) (Supplementary Table S6). In 10 out of 19 attempts, at least 80% of the restrained central helix base pairs formed. In 9 of these attempts, 100% of the restrained central helix base pairs formed. In every attempt, we observed the formation of at least 16% of these base pairs. On the other hand, strand exchange was difficult for the holo hybrid model to achieve (Figure 6E). In only 3 out of 19 attempts, 80% of the restrained central helix base pairs formed. Notably, in 6 out of 19 attempts, all the restrained base pairs stalled, and the original P1 helix remained entirely intact. The highest percentage of central helix formation achieved out of all attempts was 83%. To further confirm these results, we ran a second series of 2D REMD simulations (9 replicas, 47 ns per replica) on each hybrid model with a gradient of weak harmonic bias restraints applied to seven central helix base pairs (Supplementary Table S7). For the apo2 state, strand exchange was successful in 10 out of 19 attempts and at least 14% of restrained central helix base pairs formed in every attempt. Conversely, for the holo state, strand exchange was not successful in any of the 19 attempts. In 4 of these attempts, the restrained central helix base pairs stalled, and the P1 helix remained fully intact. The highest percentage of central helix formation achieved out of all attempts was 71%. The agreement of the 47 ns/replica and 22 ns/replica results indicates the robustness of the observed results with respect to total simulation length, as all strand exchanges sterically able to occur under the specific biases forces applied have either already happened or completely stalled within this simulation timeframe.

We also explored the experimental finding that the U91G mutant (Figure 4C) (position 80 in our holo hybrid model) yield a broken ‘ON’ phenotype. We mutated position 80 to G in our holo hybrid model to create a G:G mistmach in the P1 helix. We then ran a series of 2D REMD simulations (9 replicas, 38 ns per replica) on this mutant and again applied a gradient of weak harmonic bias restraints to seven central helix base pairs. Strand exchange was not successful in any of the 19 attempts, but partial central helix formation was observed in 12 attempts (Supplementary Table S8). This result is to be expected, as our simulations already bias central helix formation very significantly. The introduction of the U80G mutation was only sufficient to destabilize the P1 helix but did not encourage stability in the central helix. Altogether, the strand invasion 2D REMD calculations entailed ∼1.6 µs of cumulative all-atom MD simulations.

## DISCUSSION

Previous secondary structure predictions of the *yxjA* riboswitch show that the ligand binding junction of the AD and EP do not share direct overlapping sequences (43). This feature differs from the well-characterized *pbuE* purine riboswitch where a significant portion of the junction of the AD can be broken by the terminator when there is no ligand bound. Yet, in the *yxjA* riboswitch, which serves as an interesting contrasting model as an ‘OFF’ purine riboswitch, ligand binding is still able to induce a conformational change in the downstream EP, even without the junction of the AD overlapping with the EP. Our structural, functional, and simulation studies of the *yxjA* riboswitch in this work support a model in which the riboswitch relies on the formation of an intermediate central helix, mutually exclusive with both the P1 helix and the terminator helix, that mitigates the structural switching mechanism. More specifically, our data also suggests that ligand binding in the AD induces a conformational change in the P1 helix and prevents the central helix from forming, thereby allowing the terminator to form in the presence of ligand.

In either scenario, for this process to occur fast enough to respond to a bound ligand and influence whether RNAP continues to transcribe or terminate, the central helix invading the P1 helix must occur cotranscriptionally. The transition through sequential structures during transcription is a critical feature that allows transcriptional riboswitches to regulate gene expression. Our investigation of how strand exchange is involved in the mechanism of the *yxjA* riboswitch adds another example to the growing repertoire of studies showing the important role of strand exchange and cotranscriptional folding in functional RNAs, such as the rearrangement of the *E. coli* SRP RNA into a large hairpin and conformational switching of riboswitches such as ZTP, FMN and *pbuE* (28-30, 34, 42).

Our work also builds on previous studies about cotranscriptional folding and riboswitch-mediated gene regulation by combining several complementary approaches to build a comprehensive understanding of the *yxjA* riboswitch mechanism. While our previous work using cotranscriptional SHAPE-Seq has attempted to chemically probe RNA structures during transcription at all intermediate lengths during transcription (27, 29), we decided to focus our experiments in this study on a fewer number of specific single length RNAs. This approach allowed us to achieve a higher read depth for a single length when sequencing, although at the cost of losing structural information from other lengths that would have informed a matrix of reactivity values for illustrating an entire “folding pathway” of structures. However, we were still able to observe largely replicable reactivity patterns across replicates and test hypotheses by using *in vivo* functional reporter assays, experimentally informed structure modeling algorithms, and *in silico* modeling that produced results that agreed with each other.

Regarding our mechanism model, we also present how a riboswitch without a directly overlapping AD junction and EP can be influenced by ligand binding. In the case of the *yxjA* riboswitch, we have found that ligand binding creates a more stable P1 helix, which can explain how the AD “communicates” over a relatively longer distance with the downstream EP to influence its conformation. Compared to the *pbuE* riboswitch and more broadly riboswitches where the AD and EP directly overlap, this mechanism could be a central factor in determining how similar aptamers can have different expression logic depending on variations in the EP.

Several riboswitches that have been studied to date with a similar modular architecture that have no direct overlap between the AD junction and EP may involve this mechanism as well. Our results in this study suggest that a central helix motif that lies in between the terminator and the AD P1 helix may be the key feature that determines the ‘OFF’ switching expression logic. For example, a previous study by Ceres, et al. noted the modularity of select riboswitches that turn ‘OFF’ gene expression in response to ligand-binding (37). Based on their sequence complementary patterns, these riboswitches, such as SAM (*metE* and *yitJ*), lysine (*lysC*), purine (*xpt*), and FMN (*ribD*), appear to also have a P1 helix and central helix that do not overlap into the junction part of the AD, similar to the *yxjA* riboswitch. We hypothesize that these riboswitches also rely on a similar P1 helix/central helix interaction to communicate ligand binding between the AD to the EP.

Riboswitches remain an informative model for understanding how RNAs effectively fold into highly specific structures to enact their functions, especially in the context of gene regulation. They are key to understanding how cotranscriptional folding and strand exchange processes can play a reoccurring role in facilitating the folding of RNA into their specialized roles in the cell. Our observations of the *yxjA* riboswitch and its dependency on a central helix motif and strand exchange mechanism could also hint at a process central to RNA folding across biology.

## DATA AVAILABILITY

Detailed installation and usage guide for Spats is available at Read the Docs (https://spats.readthedocs.io/en/master/) Reconstructing RNA Dynamics from Data (R2D2) is available at the GitHub Repository (https://github.com/LucksLab/R2D2)

## ACCESSION NUMBERS

Sequencing data generated in this work is being deposited on the Small Read Archive (http://www.ncbi.nlm.nih.gov/sra) and will be accessible via a BioProject accession number once deposited.

SHAPE-Seq reactivity data generated in this work is being deposited on the RNA Mapping Database (http://rmdb.stanford.edu/repository/) and will be accessible using RMDB ID numbers once deposited.

## SUPPLEMENTARY DATA

Supplementary Data are available at NAR online.

## ACKNOWLEDGEMENTS

We thank Rob Batey for many helpful conversations in discussing this work, Yun-Xing Wang for discussions about aptamer-ligand interactions, Jim Brink and Steve Hockema (496code) for developing Spats v2.0.5 for SHAPE-Seq analysis, Molly Evans for feedback on manuscript drafts and improving SHAPE-Seq protocols and data analysis, and Katherine Berman for suggestions and feedback on experiments and figures.

