## Supplementary Information for "Cotranscriptional RNA strand exchange underlies the gene regulation mechanism in a purine-sensing transcriptional riboswitch"

*Corresponding authors

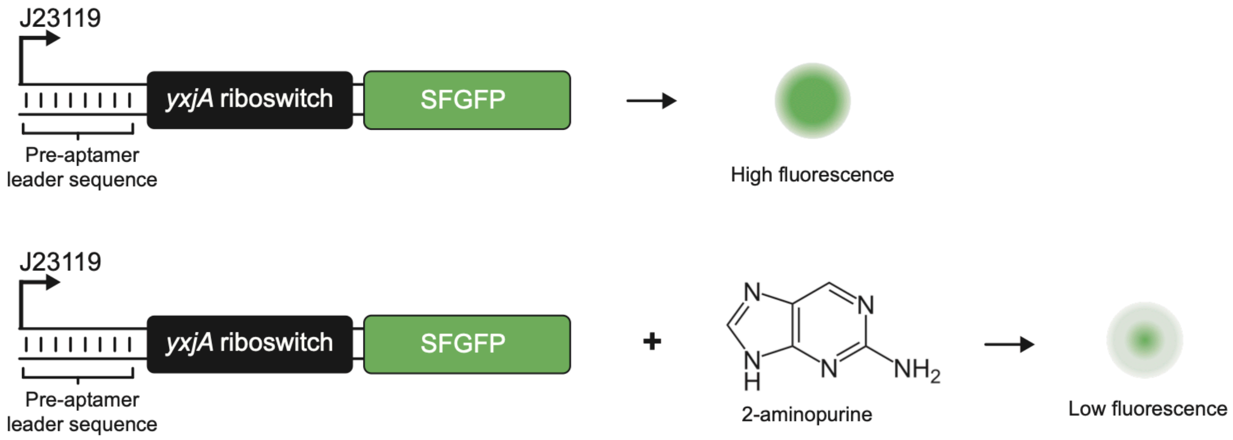


**Supplementary Figure S1 | Schematic of the *yxjA* riboswitch GFP reporter construct.** Labeled blocks represent elements included in the DNA template designed to produce super folder GFP (SFGFP) fluorescence output in response to the presence or absence of the 2-aminopurine (2AP) ligand. The DNA template consists of the *Escherichia coli* sigma 70 consensus promoter (J23119), the *Bacillus subtilis yxjA* riboswitch sequence including a 5 nt pre-aptamer leader sequence that precedes the natural riboswitch sequences, a strong synthetic ribosome binding site, and the SFGPF coding sequence. These sequences were cloned into a p15a plasmid backbone with chloramphenicol resistance using inverse polymerase chain reaction (iPCR). *E. coli* strains carrying the plasmid expressed SFGFP when grown in LB media or M9 minimal media for fluorescence assays. In the presence of a ligand such as 2-aminopurine (2AP), strains expressed relatively lower amounts of SFGFP with lower levels of fluorescence. See Supplementary Table S1 for a list of construct sequences used in this study.

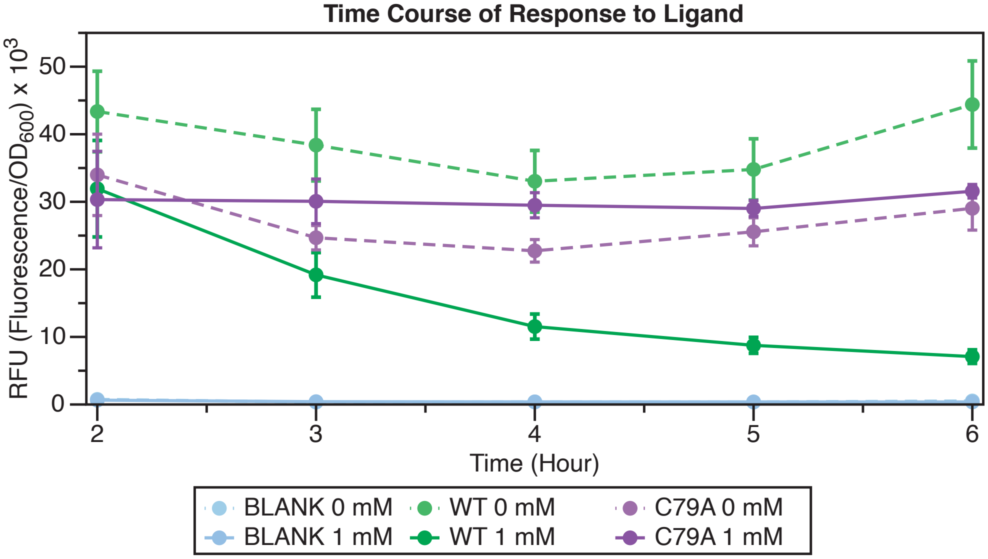


**Supplementary Figure S2 | *E. coli* GFP reporter time course optimization.** Hourly data points were collected to identify the optimal time to grow strains and measure fluorescence for subsequent experiments. Cells containing the wild type (WT) reporter plasmid, a plasmid containing the C79A ligand non-binding mutation, or a control plasmid (BLANK) lacking the riboswitch cassette, were grown overnight in LB media, split into three subcultures and then transferred to M9 minimal media for the time course either with or without the indicated ligand (0 mM or 1 mM 2AP) concentration. Bulk fluorescence was measured on a plate reader every hour between 2 and 6 hours. The 5 hour time point appeared to be optimal for observing the largest difference in fluorescence between +/- ligand conditions. At 6 hours, lower optical densities (OD) were likely to have produced artificially higher fluorescence measurements due to potentially toxic cell growth conditions. Dots represent averages from three biological replicates, each performed in triplicate technical replicates for a total of nine data points (n=9), with error bars representing standard deviation.

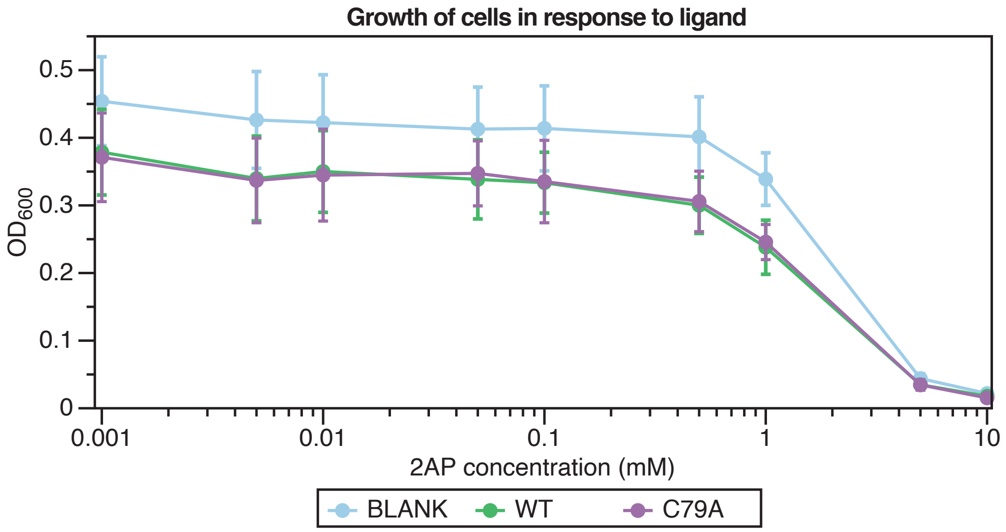


**Supplementary Figure S3 | Ligand concentration optimization.** Measurements of optical density (OD) of *E. coli* cells containing the wild type (WT) reporter plasmid, a plasmid containing the C79A ligand non-binding mutation, or a control plasmid (BLANK) lacking the riboswitch cassette, in response to different concentrations of ligand (2AP). Cells were grown overnight in LB media, split into three subcultures and then transferred to M9 minimal media with various concentrations of ligand. Bulk fluorescence was measured on a plate reader at 5 hours based on the previous time course optimization experiment (Supplementary Figure S2). Ligand concentrations higher than 1 mM, such as 5 and 10 mM showed a marked decrease in OD. Dots represent averages from three biological replicates, each performed in triplicate technical replicates for a total of nine data points (n=9), with error bars representing standard deviation.

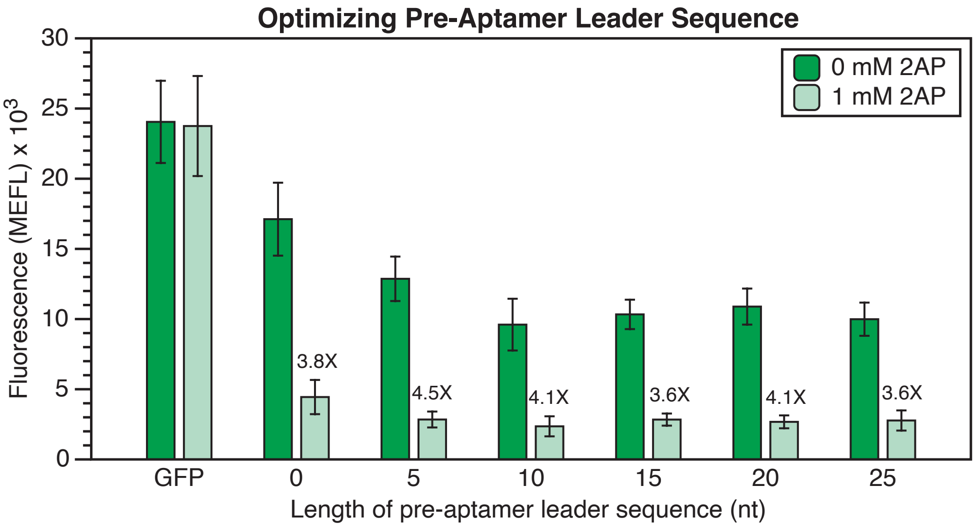


**Supplementary Figure S4 | Pre-aptamer leader sequence optimization.** GFP reporter assay data was collected from *E. coli* cells transformed with riboswitch reporter plasmids containing various pre-aptamer leader sequence lengths to identify an optimal construct. Stretches of genomic sequence located before the aptamer region of the *yxjA* riboswitch were inserted into the reporter construct between the promoter and the riboswitch aptamer domain (AD) at increments of 5 nucleotides. Inserting 5 nucleotides of the genomic pre-aptamer sequence into the reporter construct resulted in the highest fold decrease in GFP fluorescent measurement. Fold change was calculated by dividing GFP expression in the ligand condition (1 mM 2AP) by the GFP expression in the no ligand condition (0 mM 2AP) of the same construct. 'GFP' represents a control construct consisting of a constitutively expressed SFGFP preceded by a stability hairpin. Bars represent averages from three biological replicates, each performed in triplicate technical replicates for a total of nine data points (n=9), with error bars representing standard deviation.

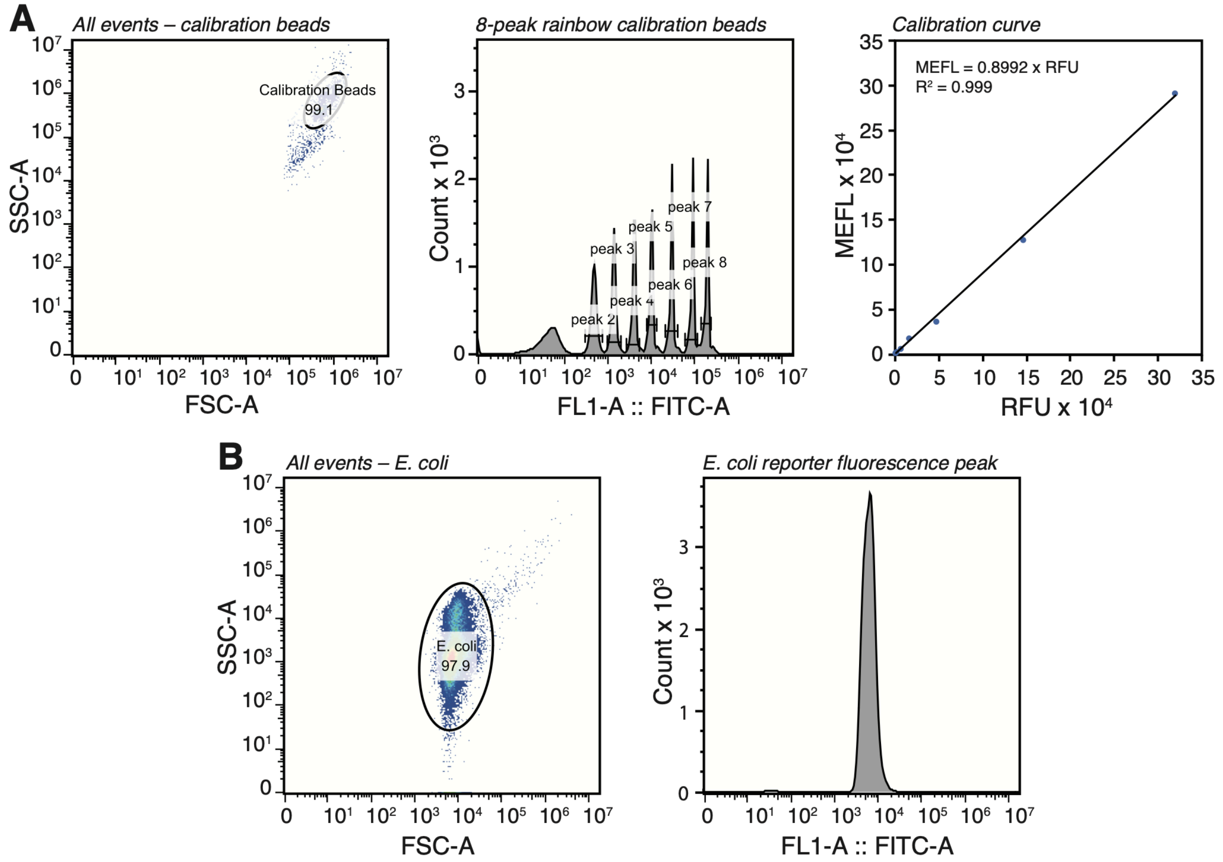


**Supplementary Figure S5 | Flow cytometry gating and calibration workflow.**
**(A)** Gating and selecting peaks to calibrate intensity measured as relative fluorescence units (RFU) into molecules of equivalent fluorescein (MEFL) with 8-peak Rainbow Particles (BD Biosciences). The bead population was selected using the FSC-A vs SSC-A profile (left). The mean fluorescence intensity (MFI) of FITC measured on the flow cytometer (BD Accuri C6 Plus) corresponding to individual peaks (center) of each bead population was plotted against MEFL values provided by the manufacturer (right). The calibration curve was calculated using a linear regression with y-intercept set to zero. The resulting multiplier value was used to convert measured RFU values into MEFL units of all *E. coli* fluorescence measurements. **(B)** Example of FSC-A vs SSC-A profile of *E. coli* populations for measuring SFGFP fluorescence using the FITC channel. A representative gate is shown on the left, and the resulting fluorescence peak of gated cells on the right. Fluorescence measurements were collected using BD CSampler software and peak averages were calculated on FlowJo 10.4.1. Histograms and plots were also generated on FlowJo 10.4.1 software. Histogram averages were converted to MEFL values using the calibration curve in (A) before being used in averages and standard deviations within individual data sets.

**A 135**

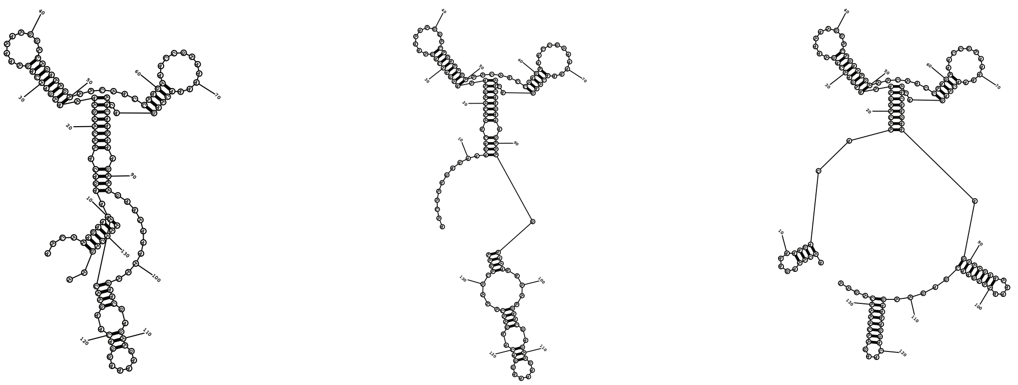

Energy = -25.6 Energy = -25.2 Energy = -24.7

**B 160**

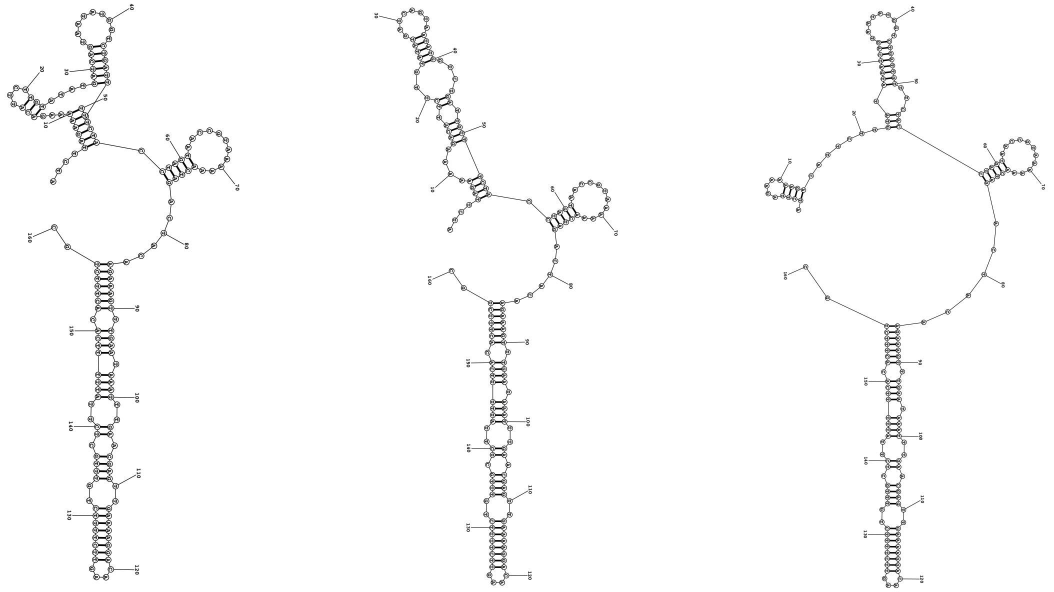

Energy = -33.8 Energy = -33.4 Energy = -32.2

**C 220**

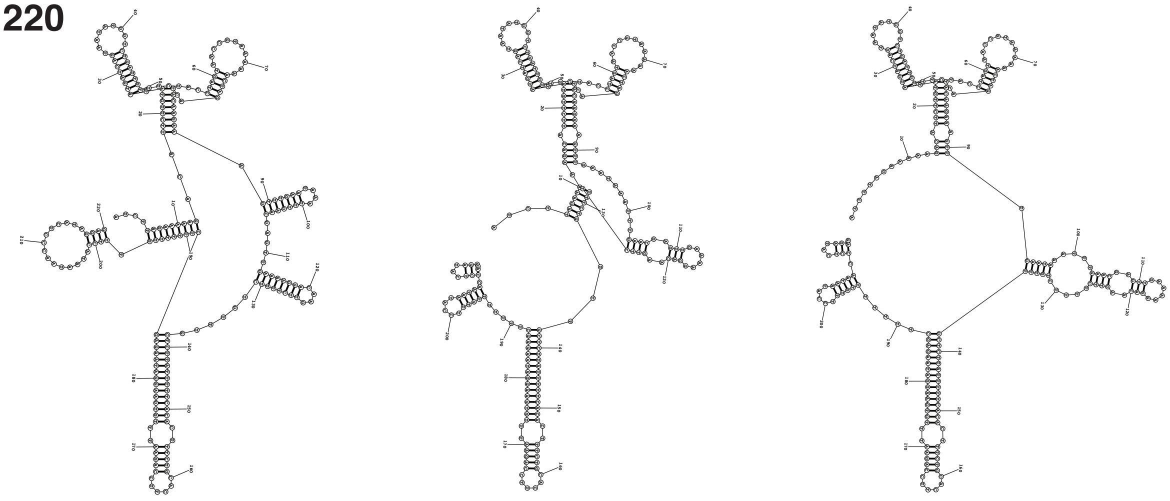

Energy = -56.1 Energy = -55.6 Energy = -55.6

**Supplementary Figure S6 | Equilibrium secondary structure predictions of *yxjA* riboswitch lengths.** Secondary structure predictions of *yxjA* riboswitch intermediate lengths and their minimum free energies generated by RNAStructure Fold (1). The minimum free energy (kcal/mol) and the first two suboptimal structures are shown. The following parameters were used for generating predictions: 10% Maximum Energy Difference; 20 Maximum Number of Structures; 3 Window Size; 30 Maximum Loop Size; 310.15 K Temperature; no SHAPE Constraints. Predictions were run for each length of the riboswitch with intermediate lengths of **(A)** 135 nt, **(B)** 160 nt, and **(C)** 220 nt (full length) riboswitch which corresponded to the DNA templates used for SHAPE-Seq experiments.

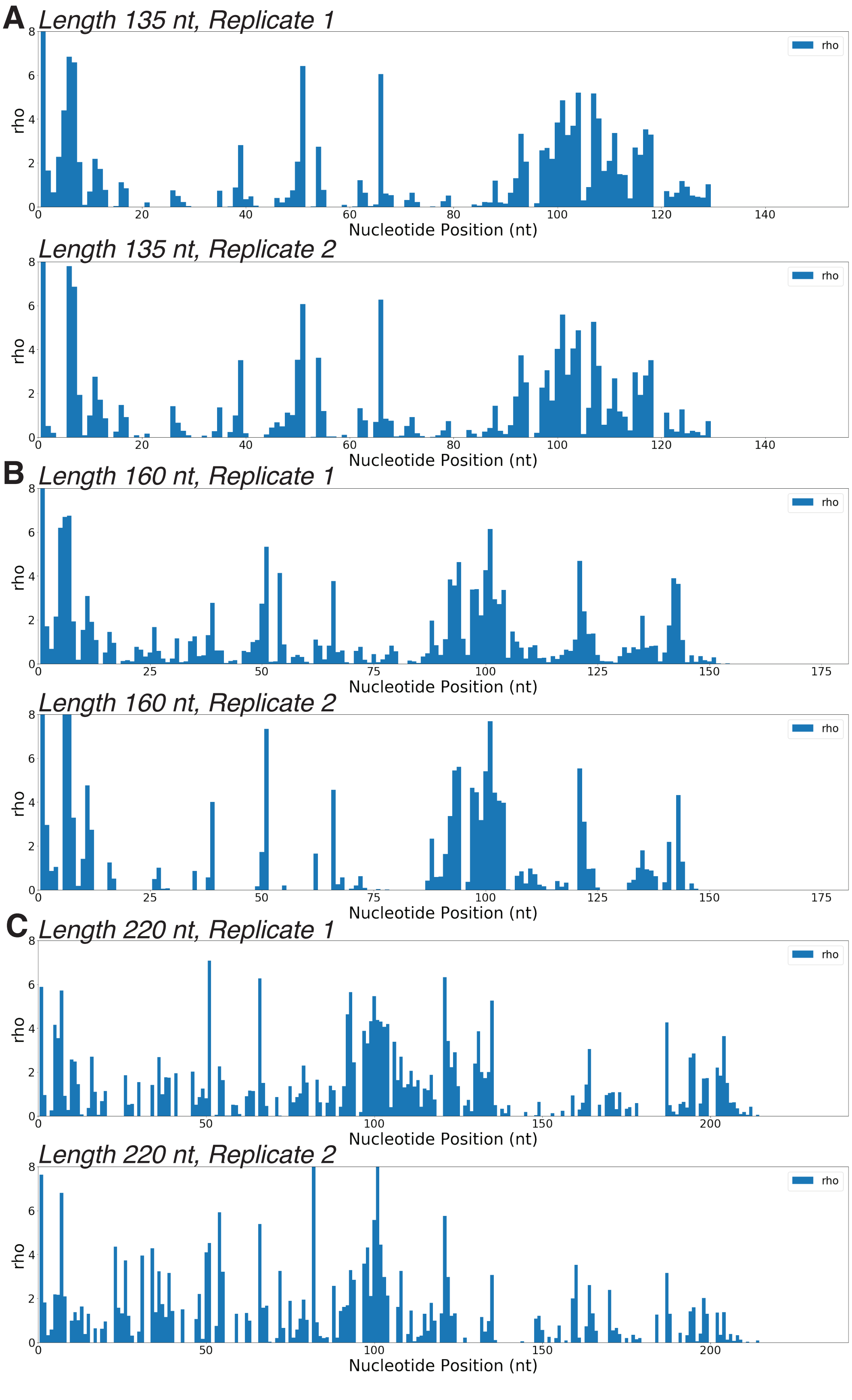
A
Length 135 nt, Replicate 1
Length 135 nt, Replicate 2
B
Length 160 nt, Replicate 1
Length 160 nt, Replicate 2
C
Length 220 nt, Replicate 1
Length 220 nt, Replicate 2
rho
Nucleotide Position (nt)

**Supplementary Figure S7 | Cotranscriptional SHAPE-Seq reactivities.** Histogram of reactivity data for several lengths of the *yxjA* riboswitch probed with benzoyl cyanide (BzCN) during *in vitro* transcription. Data includes intermediate lengths **(A)** 135 nt, (B) 160 nt, and **(C)** 220 nt (full length) riboswitch with two replicates for each. Data is reported as rho reactivities (2).

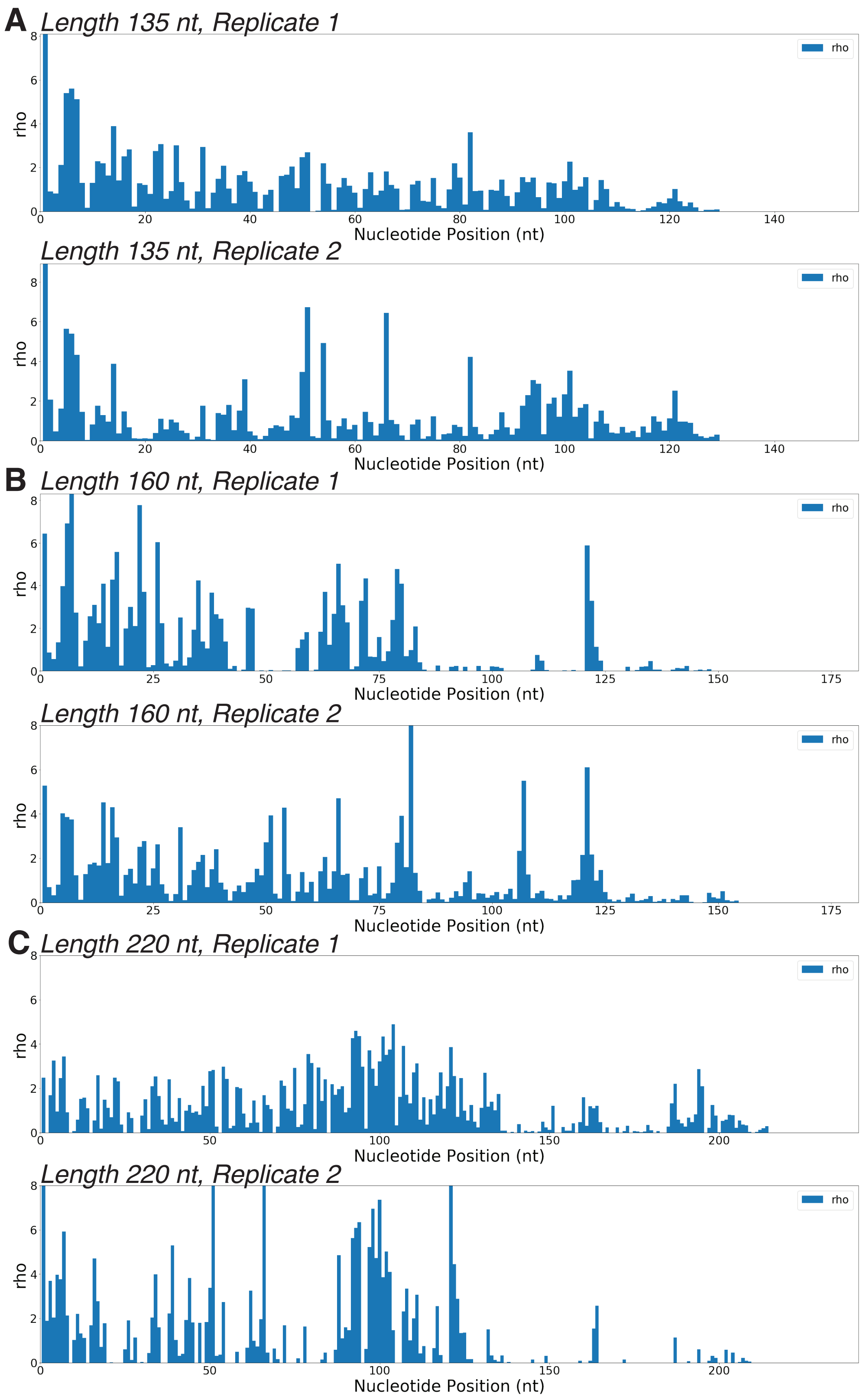
A
Length 135 nt, Replicate 1
rho
Nucleotide Position (nt)
Length 135 nt, Replicate 2
rho
Nucleotide Position (nt)
B
Length 160 nt, Replicate 1
rho
Nucleotide Position (nt)
Length 160 nt, Replicate 2
rho
Nucleotide Position (nt)
C
Length 220 nt, Replicate 1
rho
Nucleotide Position (nt)
Length 220 nt, Replicate 2
rho
Nucleotide Position (nt)

**Supplementary Figure S8 | Equilibrium SHAPE-Seq reactivities.** Histogram of reactivity data for several lengths of the *yxjA* riboswitch probed with BzCN once denatured and refolded after *in vitro* transcription (see Methods). Data includes intermediate lengths **(A)** 135 nt, **(B)** 160 nt and **(C)** 220 nt (full length) riboswitch with two replicates for each. Data is reported as rho reactivities (2).

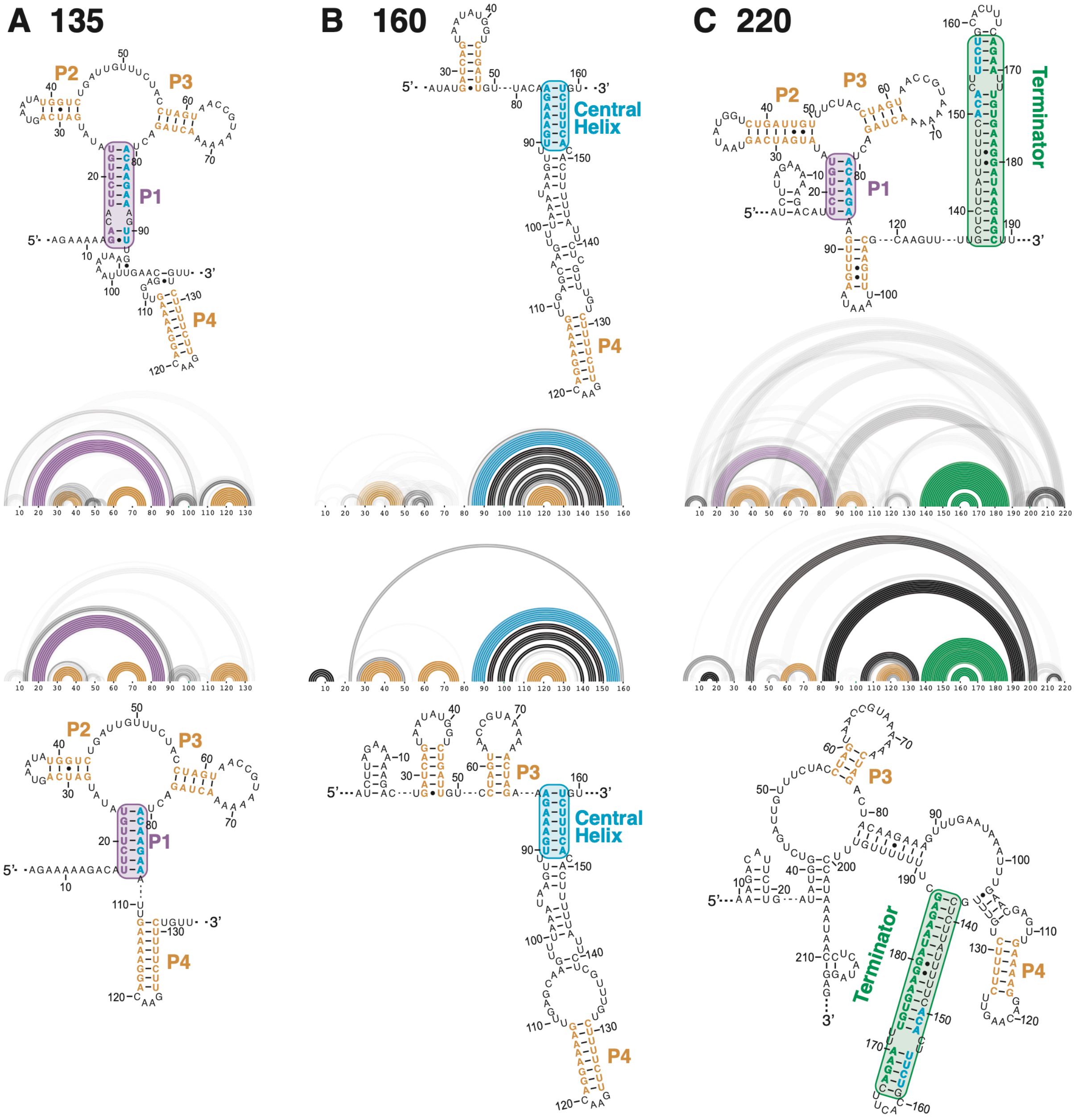


**Supplementary Figure S9 | Experimentally informed equilibrium secondary structure models of *yxjA* riboswitch lengths.** Secondary structure models generated by R2D2 (3) sing equilibrium SHAPE-Seq data of single-lengths of the *yxjA* riboswitch without ligand. Equilibrium SHAPE-Seq experiments were performed at two intermediate lengths (**(A)** 135 nt, **(B)** 160 nt) and for the full length riboswitch (**(C)** 220 nt ) in the absence of 2AP. Reactivity data were then incorporated into R2D2 pathway modeling to generate one hundred structures that are maximally consistent with the reactivity data. This family of structures is displayed as RNAbow plots, where an arc between two positions indicates a base pair in a specific structure, and the opacity of the arc indicates the prevalence of the base pair among the selected structures. Two replicates were performed at each length. The consensus secondary structure over the population of selected structures is shown as a secondary structure above or below each bow plot. In the secondary structures, the central helix is colored teal while the 5′ side of P1 is purple and the 3′ side of the terminator is colored green. Colored boxes drawn around the P1 helix, central helix, and terminator match the color of arcs in the RNAbow plots.

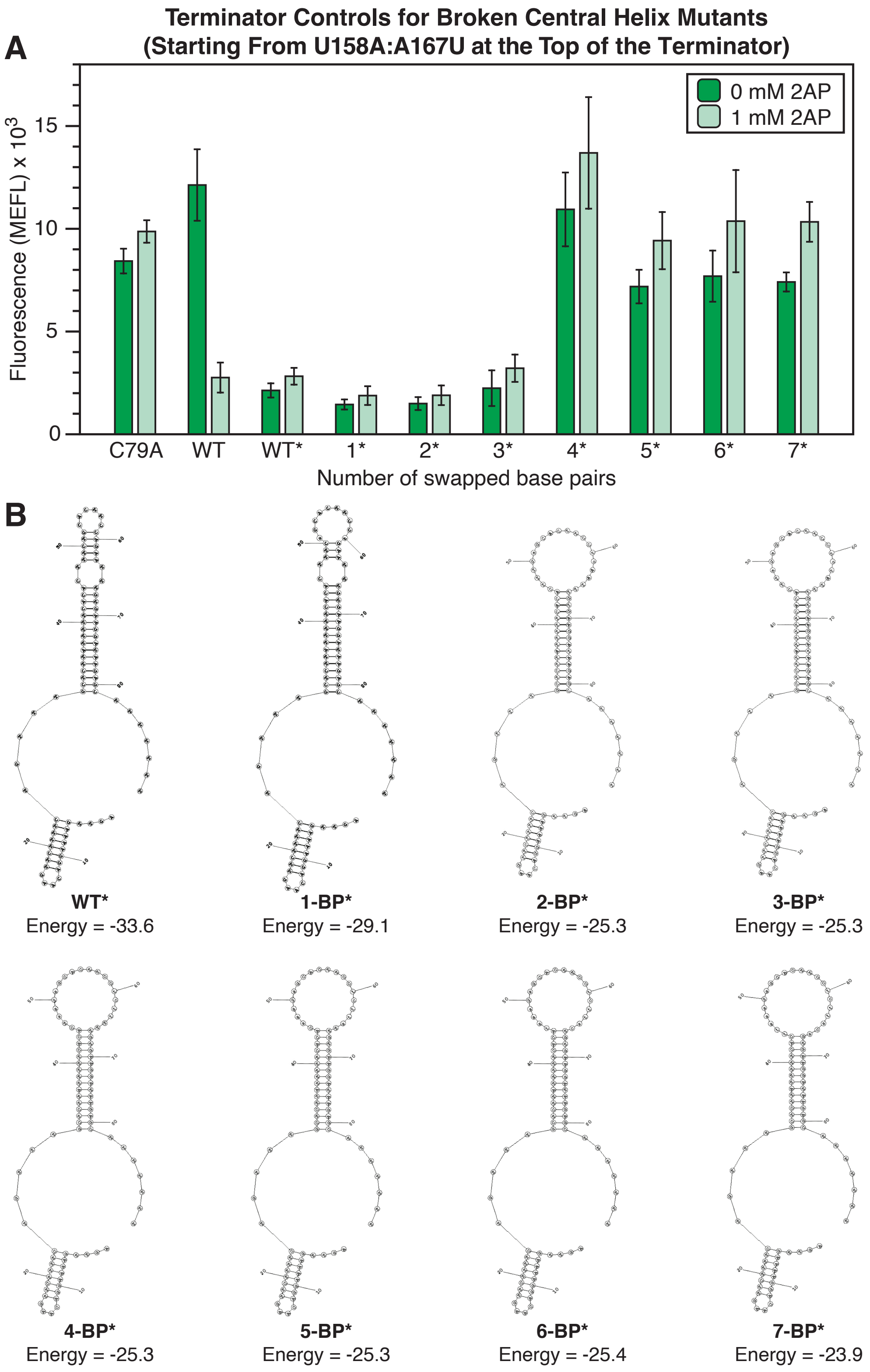
A
Terminator Controls for Broken Central Helix Mutants
(Starting From U158A:A167U at the Top of the Terminator)
0 mM 2AP
1 mM 2AP
Fluorescence (MEFL) x 10^3
0
5
10
15
C79A
WT
WT*
1*
2*
3*
4*
5*
6*
7*
Number of swapped base pairs
B
WT*
Energy = -33.6
1-BP*
Energy = -29.1
2-BP*
Energy = -25.3
3-BP*
Energy = -25.3
4-BP*
Energy = -25.3
5-BP*
Energy = -25.3
6-BP*
Energy = -25.4
7-BP*
Energy = -23.9

**Supplementary Figure S10 | Terminator controls for broken central helix mutants with mutations introduced from U158A:A167U towards A152U:U173A (top of terminator towards bottom of terminator).** *E. coli* cells were transformed with either riboswitch expression constructs, or control constructs that contain a promoter followed by a terminator sequence, a ribosome binding site (RBS), and the SFGFP coding sequence. Terminator sequences contained the same mutations in the terminator as the corresponding central helix mutants in Figure 5. Terminator constructs were generated from the sequences in Figure 5 by deleting the aptamer domain sequence (nts 12 – 91). **(A)** Fluorescence data of GFP reporter assays collected with flow cytometry with and without ligand (1 mM 2AP) added to subcultures. C79A represents the full riboswitch construct with the ligand non-binding mutation introduced, WT represents the wild-type full riboswitch expression construct, WT* represents the WT terminator construct, and numbers indicate the number of base pairs swapped in the terminator expression construct starting from the mutation U158A:A167U and going down the terminator to the A152U:U173A mutation. **(B)** Equilibrium secondary structure predictions and minimum free energies (kcal/mol) of the terminator sequences generated by RNAStructure (1) with the same settings as Supplementary Figure 6. Bars represent averages from three biological replicates, each performed in triplicate technical replicates for a total of nine data points (n=9), with error bars representing standard deviation.

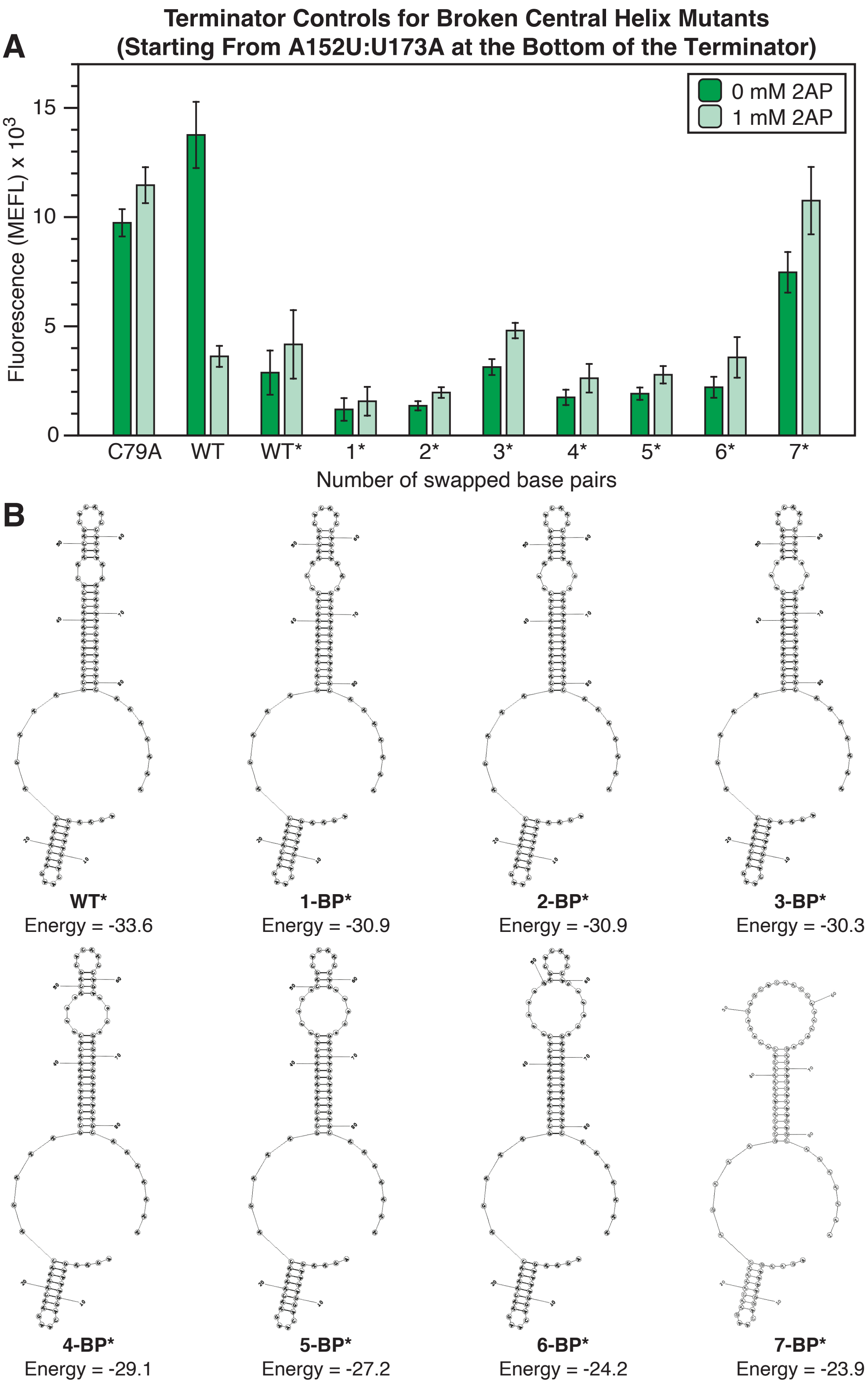

A
Terminator Controls for Broken Central Helix Mutants
(Starting From A152U:U173A at the Bottom of the Terminator)
0 mM 2AP
1 mM 2AP
Fluorescence (MEFL) x 10^3
0
5
10
15
C79A
WT
WT*
1*
2*
3*
4*
5*
6*
7*
Number of swapped base pairs
B
WT*
Energy = -33.6
1-BP*
Energy = -30.9
2-BP*
Energy = -30.9
3-BP*
Energy = -30.3
4-BP*
Energy = -29.1
5-BP*
Energy = -27.2
6-BP*
Energy = -24.2
7-BP*
Energy = -23.9

**Supplementary Figure S11 | Terminator controls for broken central helix mutants with mutations introduced from the A152U:U173A towards U158A:A167U (bottom of terminator towards top of terminator).** *E. coli* cells were transformed with either riboswitch expression constructs, or control constructs that contain a promoter followed by a terminator sequence, a ribosome binding site (RBS), and the SFGFP coding sequence. Terminator sequences contained the same mutations in the terminator as the corresponding central helix mutants in Figure 5. Terminator constructs were generated from the sequences in Figure 5 by deleting the aptamer domain sequence (nts 12 – 91). **(A)** Fluorescence data of GFP reporter assays collected with flow cytometry with and without ligand (1 mM 2AP) added to subcultures. C79A represents the full riboswitch construct with the ligand non-binding mutation introduced, WT represents the wild-type full riboswitch expression construct, WT* represents the WT terminator construct, and numbers indicate the number of base pairs swapped in the terminator expression construct starting from the mutation A152U:U173A and going up the terminator to the U158A:A167U mutation. **(B)** Equilibrium secondary structure predictions and minimum free energies (kcal/mol) of the terminator sequences generated by RNAStructure (1) with the same settings as Supplementary Figure 6. Bars represent averages from three biological replicates, each performed in triplicate technical replicates for a total of nine data points (n=9), with error bars representing standard deviation.

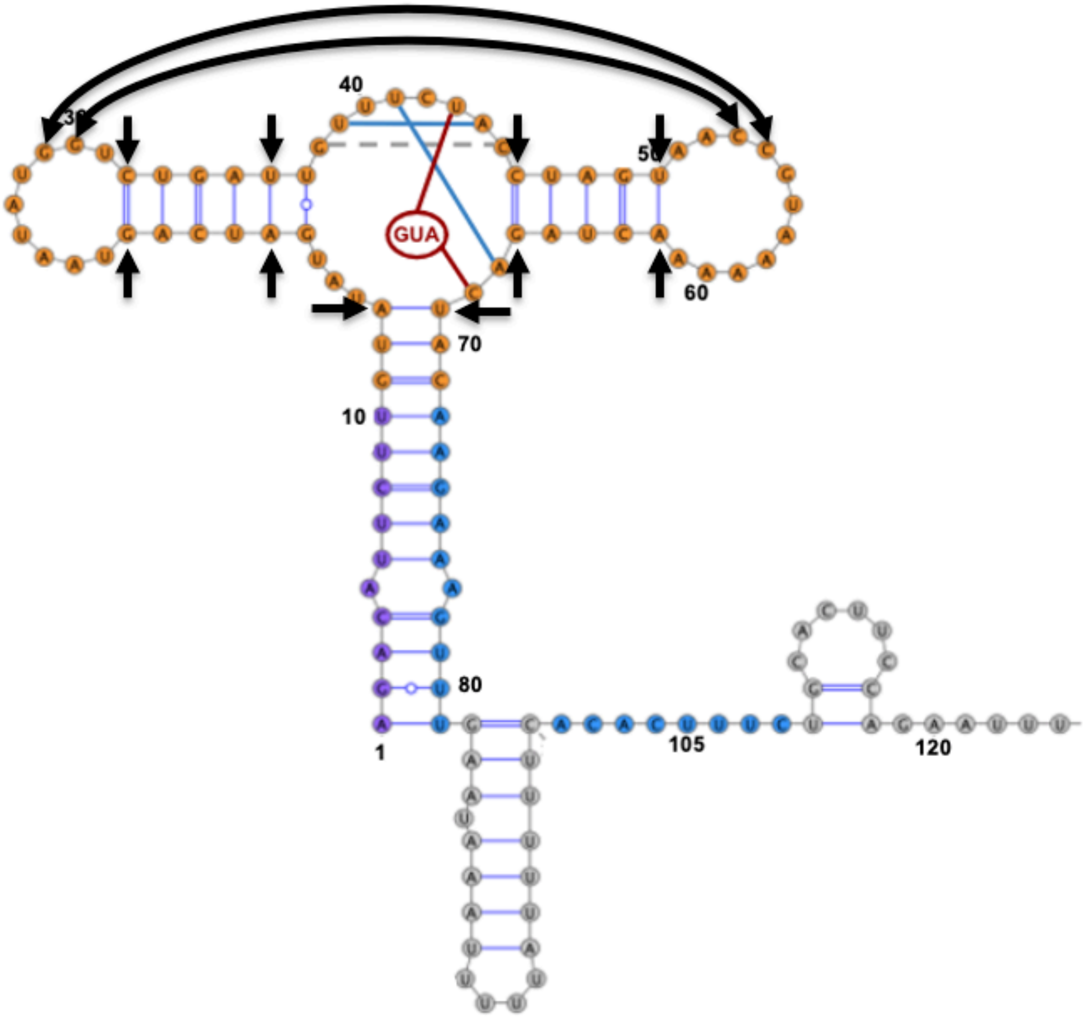


**Supplementary Figure S12 | Base pairs in the aptamer that were restrained during the equilibration of the apo1, apo2 and holo hybrid models.** Harmonic bias restraints were applied to the designated base pairs (black arrows) at a strength of 5 kcal/mol during the equilibration phase of molecular dynamics (MD) simulations run on the apo1, apo2 and holo hybrid models. Blue lines connecting nucleotides in the three-way junction (3WJ) of the aptamer represent harmonic bias restraints applied to base pairing interactions in the apo state while red lines represent restraints applied to base pairing interactions between bound guanine and the 3WJ in the holo state.

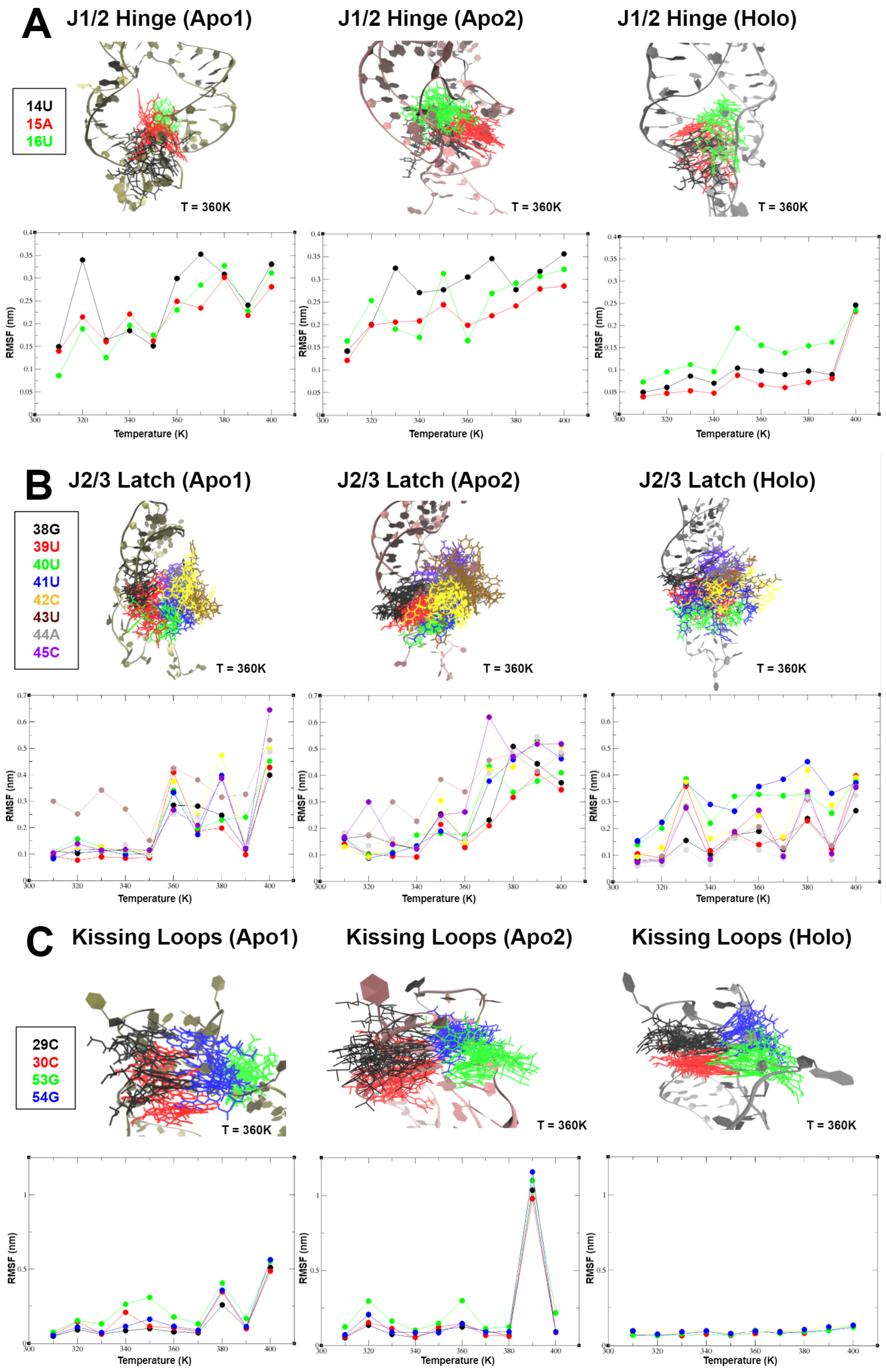
A
J1/2 Hinge (Apo1)
J1/2 Hinge (Apo2)
J1/2 Hinge (Holo)
14U
15A
16U
T = 360K
RMSF (nm)
Temperature (K)
B
J2/3 Latch (Apo1)
J2/3 Latch (Apo2)
J2/3 Latch (Holo)
38G
39U
40U
41U
42C
43U
44A
45C
C
Kissing Loops (Apo1)
Kissing Loops (Apo2)
Kissing Loops (Holo)
29C
30C
53G
54G

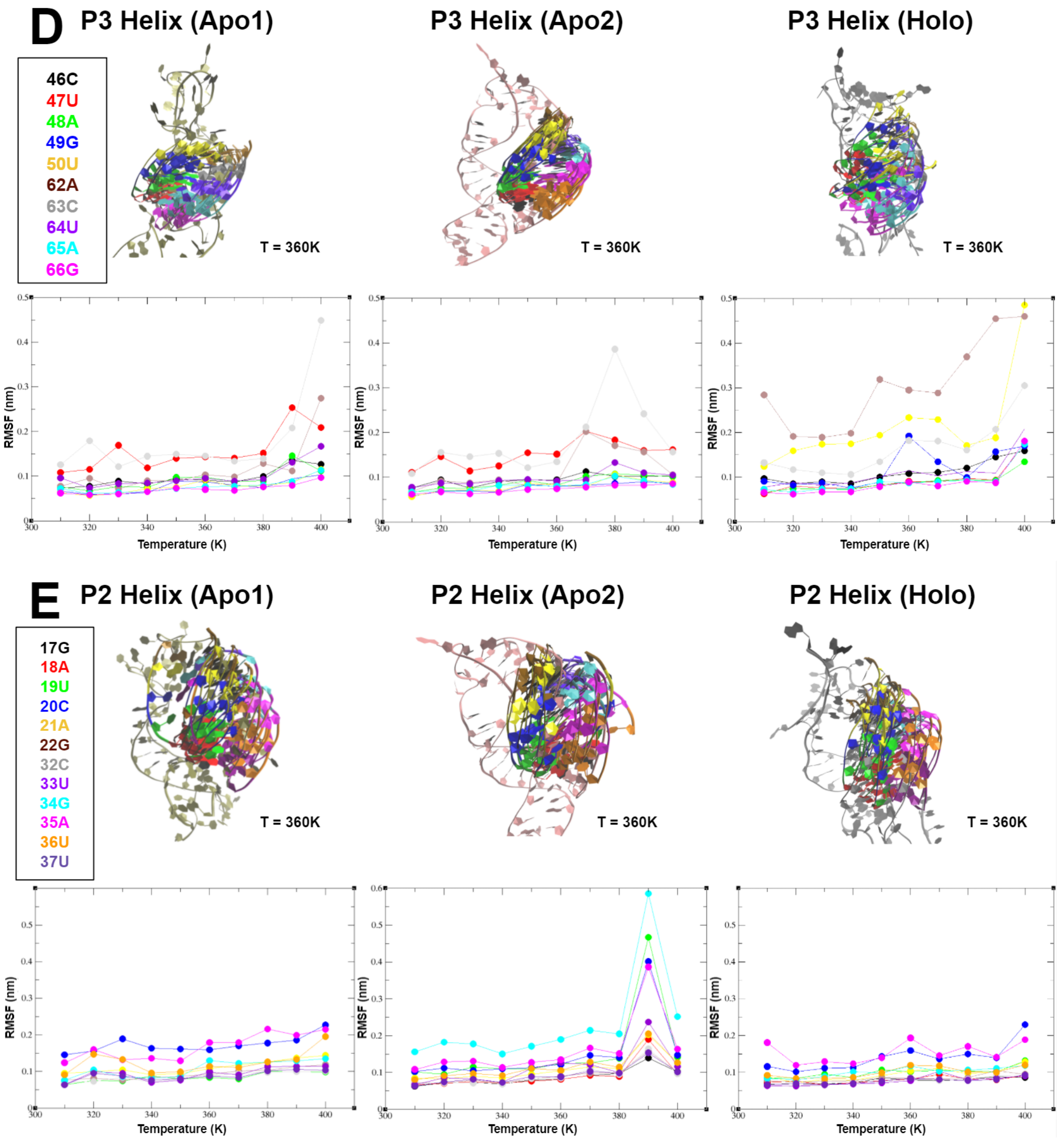


**Supplementary Figure S13 | Per-residue root mean square fluctuation (RMSF) analysis of the *yxjA* aptamer.** Per-residue RMSF was calculated over the course of a 50 ns MD simulation for each of five tertiary structural elements within the apo1, apo2 and holo aptamer models: **(A)** J1/2 hinge, **(B)** J2/3 latch, **(C)** kissing loops, **(D)** P3 helix, and **(E)** P2 helix. Nucleotide residues that were analyzed are indicated in the legend for each element. Simulations were performed at a range of temperatures (310 K – 400 K) and RMSF was calculated as the average deviation between the positions of a residue over the course of a simulation and its reference position. Snapshots of structures are shown above each plot for T = 360 K.

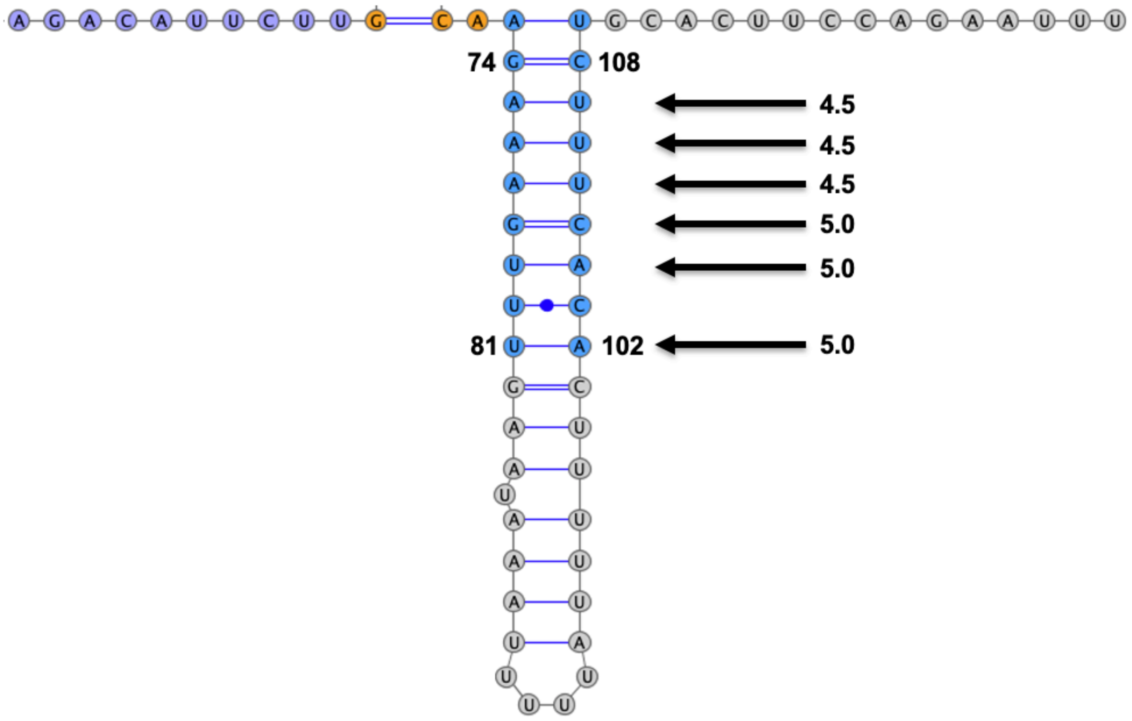


**Supplementary Figure S14 | Gradient of harmonic bias restraints applied to base pairs involved in strand exchange.** A gradient of harmonic bias restraints (black arrows) was applied to nudge the apo2 and holo hybrid models towards the anti-terminated (central helix formed) conformation. The strength of each restraint is indicated by numbers beside each arrow (kcal/mol). The strongest restraints were applied at the bottom of the P4 helix, while the weakest restraints were applied at the top to encourage strand exchange to happen naturally and within a time scale that was computationally feasible to observe.

**Supplementary Table S1 | Plasmid sequences used for *in vivo* fluorescence measurements.** Sequences for reporter plasmids transformed into *E. coli* for gene expression assays using a plate reader or flow cytometer can be found listed in the supporting data Excel file Cheng_yxjA_Strand_Exchange_Supp_Data_File1_Sequences.xlsx

**Supplementary Table S2 | Template sequences used for *in vitro* transcription.** DNA template sequences used for cotranscriptional and equilibrium SHAPE-Seq experiments are listed below. Orange represents the J23119 (SpeI) promoter sequence, teal represents the *yxjA* riboswitch sequence, green represents the RNAP footprint (14 nt), and purple represents the EcoRI restriction site. Templates for equilibrium SHAPE-Seq experiments were shifted by 14 nt in the 5′ direction to account for the lack of an RNAP footprint and match the probed structure with the corresponding cotranscriptional experiment of the same length of the riboswitch.

| RNA | Experiment | Sequence |
|---|---|---|
| *yxjA* riboswitch, C79T d5, 135 nt + RNAP footprint | Cotranscriptional | ctaaagatctttgacagctagctcagtcctaggtataatactagtATCTT AGAAAAAGACATTCTTGTATATGATCAGTAATATGG TCTGATTGTTTCTACCTAGTAACCGTAAAAAACTAG AtTACAAGAAAGTTTGAATAAATTTGAACGAGTTGA AAAGGACAAGTTCTTTTCTGTTTGCTCTTATTTTTC gaattcaaaaaaa |
| *yxjA* riboswitch, C79T d5, 160 nt + RNAP footprint | Cotranscriptional | ctaaagatctttgacagctagctcagtcctaggtataatactagtATCTT AGAAAAAGACATTCTTGTATATGATCAGTAATATGG TCTGATTGTTTCTACCTAGTAACCGTAAAAAACTAG AtTACAAGAAAGTTTGAATAAATTTGAACGAGTTGA AAAGGACAAGTTCTTTTCTGTTTGCTCTTATTTTTC ACACTTTCTGCACTTCCAGAATTTGgaattcaaaaaaa |
| *yxjA* riboswitch, C79T d5, full length | Cotranscriptional, equilibrium | ctaaagatctttgacagctagctcagtcctaggtataatactagtATCTT AGAAAAAGACATTCTTGTATATGATCAGTAATATGG TCTGATTGTTTCTACCTAGTAACCGTAAAAAACTAG AtTACAAGAAAGTTTGAATAAATTTGAACGAGTTGA AAAGGACAAGTTCTTTTCTGTTTGCTCTTATTTTTC ACACTTTCTGCACTTCCAGAATTTGTGAAGGATAA GAGCTTTTTTTGTTTCCATAATAACCCTCATAGGAG |
| *yxjA* riboswitch, C79T d5, 135 nt | Equilibrium | ctaaagatctttgacagctagctcagtcctaggtataatactagtATCTT AGAAAAAGACATTCTTGTATATGATCAGTAATATGG TCTGATTGTTTCTACCTAGTAACCGTAAAAAACTAG AtTACAAGAAAGTTTGAATAAATTTGAACGAGTTGA AAAGGACAAGTTCTTTTCTGTT |
| *yxjA* riboswitch, C79T d5, 160 nt | Equilibrium | ctaaagatctttgacagctagctcagtcctaggtataatactagtATCTT AGAAAAAGACATTCTTGTATATGATCAGTAATATGG TCTGATTGTTTCTACCTAGTAACCGTAAAAAACTAG AtTACAAGAAAGTTTGAATAAATTTGAACGAGTTGA AAAGGACAAGTTCTTTTCTGTTTGCTCTTATTTTTC ACACTTTCTGC |

**Supplementary Table S3 | Oligonucleotide sequences used for SHAPE-Seq.** Oligonucleotide sequences used for cotranscriptional and equilibrium SHAPE-Seq experiments are listed below. 'XXXXXX' in the Illumina reverse primers represent TruSeq indexes. Abbreviations used are according to Integrated DNA Technologies (IDT) ordering notation.

| Description | Sequence |
|---|---|
| RNA linker | /5Phos/CUGACUCGGGCACCAAGGA/3ddC/ |
| Reverse transcription primer | /5Biosg/GTCCTTGGTGCCCGAGT |
| DNA adapter/dumbbell† | /5Phos/TGAAGAGCCTAGTCGCTGTTCANNNNNNCTGCCC/3SpC3/ |
| PE_F* | AATGATACGGCGACCACCGAGATCTACACTCTTTCCCTACACGACGCTCTTCCGATCT |
| Selection primer (+) | CTTTCCCTACACGACGCTCTTCCGATCTRRRYGCATCCACAATAGAAGAAGGATGC*C*G*C*A |
| Selection primer (-) | CTTTCCCTACACGACGCTCTTCCGATCTYYYRGCATCCACAATAGAAGAAGGATGC*C*G*C*A |
| Illumina reverse primers (TruSeq)* | CAAGCAGAAGACGGCATACGAGATXXXXXXGTGACTGGAGTTCAGACGTGTGCTCTTCCGATCTTGAACAGCGACTAGGCTCTTCA |
| Forward template amplification primer | CTAAAGATCTTTGACAGCTAGCTCAGTCCTAGGTATAATACTAGT |
| Reverse template primer for *yxjA* riboswitch 135 nt (cotranscriptional) | TTTTTTTGAATTCGAAAAATAAGAGCAAACAGAAAAGAAC |
| Reverse template primer for *yxjA* riboswitch 160 nt (cotranscriptional) | TTTTTTTGAATTCCAAATTCTGGAAGTGCAGAAAGTGTG |
| Reverse template primer for *yxjA* riboswitch full length (cotranscriptional and EQ) | CTCCTATGAGGGTTATTATGGAAACAAAAAAAGCTC |
| Reverse template primer for *yxjA* riboswitch 135 nt (EQ) | AACAGAAAAGAACTTGTCCTTTTCAACTCGTTC |
| Reverse template primer for *yxjA* riboswitch 160 nt (EQ) | GCAGAAAGTGTGAAAAATAAGAGCAAACAGAAAAG |

†Ritchey et al., *Nucleic Acids Research*, 2017 (4)

*Oligonucleotide sequences © 2007-2013 Illumina, Inc. All rights reserved.

**Supplementary Table S4 | Small Read Archive (SRA) deposition table.** Sequencing data generated in this work is being deposited on the Small Read Archive (http://www.ncbi.nlm.nih.gov/sra) and will be accessible via a BioProject accession number once deposited.

**Supplementary Table S5 | RNA Mapping Database (RMDB) deposition table.** SHAPE-Seq reactivity data generated in this work is being deposited on the RNA Mapping Database (http://rmdb.stanford.edu/repository/) and will be accessible using RMDB ID numbers once deposited.

**Supplementary Table S6 | Strand exchange statistics for the first series of two-dimensional replica exchange molecular dynamics (2D REMD) simulations.** Progression of strand exchange at the end of each simulation is described by the number of exchanged base pairs formed by the invading central helix, the percent of base pairs formed by the central helix out of the total possible base pairs the central helix could form, and whether the strand exchange attempt by the central helix was successful or not. These simulations were run for 22 ns in which the gradient of weak harmonic bias restraints was applied to six central helix base pairs (Supplementary Figure S14). Each row represents a different replica and nine replicas were included for each simulation.

| **Apo2** | | | **Holo** | | |
|---|---|---|---|---|---|
| Attempt 1 | | | Attempt 1 | | |
| *Exchanged BPs* | *% Formed* | *Success* | *Exchanged BPs* | *% Formed* | *Success* |
| 1 | 16.67 | N | 0 | 0.00 | N |
| 1 | 16.67 | N | 0 | 0.00 | N |
| 6 | 100.00 | Y | 4 | 66.67 | N |
| 6 | 100.00 | Y | 5 | 83.33 | Y |
| 6 | 100.00 | Y | 0 | 0.00 | N |
| 3 | 50.00 | N | 0 | 0.00 | N |
| 3 | 50.00 | N | 1 | 16.67 | N |
| 1 | 16.67 | N | 1 | 16.67 | N |
| 6 | 100.00 | Y | 3 | 50.00 | N |
| Attempt 2 | | | Attempt 2 | | |
| *Exchanged BPs* | *% Formed* | *Success* | *Exchanged BPs* | *% Formed* | *Success* |
| 2 | 33.33 | N | 3 | 50.00 | N |
| 5 | 83.33 | Y | 3 | 50.00 | N |
| 4 | 66.67 | N | 4 | 66.67 | N |
| 6 | 100.00 | Y | 4 | 66.67 | N |
| 6 | 100.00 | Y | 5 | 83.33 | Y |
| 6 | 100.00 | Y | 5 | 83.33 | Y |
| 6 | 100.00 | Y | 0 | 0.00 | N |
| 6 | 100.00 | Y | 0 | 0.00 | N |
| 0 | 0.00 | N | 2 | 33.33 | N |

**Supplementary Table S7 | Strand exchange statistics for the second series of two-dimensional replica exchange molecular dynamics (2D REMD) simulations.** Progression of strand exchange at the end of each simulation is described by the number of exchanged base pairs formed by the invading central helix, the percent of base pairs formed by the central helix out of the total possible base pairs the central helix could form, and whether the strand exchange attempt by the central helix was successful or not. These simulations were run for 47 ns in which the gradient of weak harmonic bias restraints was applied to seven central helix base pairs. Each row represents a different replica and nine replicas were included for each simulation.

| **Apo2** | | | **Holo** | | |
|---|---|---|---|---|---|
| Attempt 1 | | | Attempt 1 | | |
| *Exchanged BPs* | *% Formed* | *Success* | *Exchanged BPs* | *% Formed* | *Success* |
| 4 | 57.14 | N | 3 | 42.86 | N |
| 6 | 85.71 | Y | 5 | 71.43 | N |
| 4 | 57.14 | N | 4 | 57.14 | N |
| 7 | 100.00 | Y | 3 | 42.86 | N |
| 7 | 100.00 | Y | 2 | 28.57 | N |
| 3 | 42.86 | N | 0 | 0.00 | N |
| 5 | 71.43 | N | 2 | 28.57 | N |
| 6 | 85.71 | Y | 2 | 28.57 | N |
| 7 | 100.00 | Y | 2 | 28.57 | N |
| Attempt 2 | | | Attempt 2 | | |
| *Exchanged BPs* | *% Formed* | *Success* | *Exchanged BPs* | *% Formed* | *Success* |
| 7 | 100.00 | Y | 0 | 0.00 | N |
| 1 | 14.29 | N | 4 | 57.14 | N |
| 6 | 85.71 | Y | 0 | 0.00 | N |
| 7 | 100.00 | Y | 0 | 0.00 | N |
| 7 | 100.00 | Y | 4 | 57.14 | N |
| 1 | 14.29 | N | 2 | 28.57 | N |
| 6 | 85.71 | Y | 4 | 57.14 | N |
| 1 | 14.29 | N | 4 | 57.14 | N |
| 3 | 42.86 | N | 4 | 57.14 | N |

**Supplementary Table S8 | Strand exchange statistics for two-dimensional replica exchange molecular dynamics (2D REMD) simulations of the broken ON holo mutant.** Progression of strand exchange at the end of each simulation is described by number of exchanged base pairs formed by the invading central helix, the percent of base pairs formed by the central helix out of the total possible base pairs the central helix could form, and whether the strand exchange attempt by the central helix was successful or not. These simulations were run for 38 ns in which the gradient of weak harmonic bias restraints was applied to seven central helix base pairs. Each row represents a different replica, and there was a total of nine replicas per simulation.

| **Holo (U80G)** | | | **Holo (U80G)** | | |
|---|---|---|---|---|---|
| Attempt 1 | | | Attempt 2 | | |
| *Exchanged BPs* | *% Formed* | *Success* | *Exchanged BPs* | *% Formed* | *Success* |
| 0 | 0 | N | 0 | 0.00 | N |
| 2 | 28.57 | N | 3 | 42.86 | N |
| 1 | 14.29 | N | 1 | 14.29 | N |
| 3 | 42.86 | N | 2 | 28.57 | N |
| 4 | 57.14 | N | 3 | 42.86 | N |
| 0 | 0.00 | N | 2 | 28.57 | N |
| 1 | 14.29 | N | 0 | 0.00 | N |
| 0 | 0.00 | N | 1 | 14.29 | N |
| 0 | 0.00 | N | 0 | 0.00 | N |

**Supplementary Note S1 | SHAPE-Seq Data Analysis.**

### Spats:

#### Download and Installation:

Software for Spats can be downloaded from GitHub:
http://luckslab.github.io/spats/index.html
Detailed installation and documentation for the use of Spats can be found here:
https://spats.readthedocs.io/en/master/

#### Usage:

The following parameters were specified when running Spats to analyze SHAPE-Seq sequencing data reported in this work:
ignore_stops_with_mismatched_overlap = *False*
allowed_dumbbell_errors = *3*
allow_indeterminate = *True*
minimum_target_match_length = *26*
single_target_linker = *GTCCTTGGTGCCCGAGTCAG*
allow_multiple_rt_starts = *True*
count_mutations = *True*
count_edge_mutations = *"stop_and_mut"*
allowed_target_errors = *2*
mutations_require_quality_score = *30*
dumbbell = *TGAACAGCGACTAGGCTCTTCA*
count_left_prefixes = *True*
collapse_left_prefixes = *True*

## R2D2:

#### Download and Installation:

R2D2 can be downloaded from GitHub to a Linux server:
https://github.com/LucksLab/R2D2

#### Usage:

Input files are SHAPE-Seq reactivities generated by Spats 2.0.5 using the command:
spats_tool dump old_txt

The following parameters were used for cotranscriptional SHAPE-Seq data analyzed by Spats reported in this work: python <installation_dir>/R2D2/analyze_cotrans_SHAPE-Seq.py --in_dir <reactivity_dir> --out_dir <output_dir> --adapter

"CTGACTCGGGCACCAAGG" --e 50000 --endcut 0 --constrained_c "3.5" --scale_rho_max "1" --draw_all "True" -- most_count_tie_break "False" --weight_paired "0.8" --scaling_func "K" --cap_rhos "True" --pol_fp "14" --p 1

The following parameters were used for equilibrium-refolded SHAPE-Seq data analyzed by Spats reported in this work: python <installation_dir>/R2D2 /analyze_cotrans_SHAPE-Seq.py --in_dir <reactivity_dir> --out_dir <output_dir> --adapter "CTGACTCGGGCACCAAGG" --e 50000 --endcut 0 --constrained_c "3.5" --scale_rho_max "1" --draw_all "True" -- most_count_tie_break "False" --weight_paired "0.8" --scaling_func "K" --cap_rhos "True" --pol_fp "0" --p 1

**Supporting Data File Descriptions**

Cheng_yxjA_Strand_Exchange_Supp_Data_File1_Sequences.xlsx – DNA sequences used in this study.

Cheng_yxjA_Strand_Exchange_Supp_Data_File2_All_Source_Data.xlsx – Calibrated flow cytometry data for all figures.

White_yxjA_Strand_Exhange_Supp_Data_Movie1.mp4 – Molecular dynamics trajectories showing the differences in flexibility between the apo1, apo2 and holo aptamer domains.

White_yxjA_Strand_Exhange_Supp_Data_Movie2.mp4 – 2D REMD trajectories showing successful strand exchange in apo2, stalled strand exchange in holo, and partially reconstituted strand exchange in the broken 'ON' holo mutant. Colors are as in Figure 6.
